# Local B-cell immunity and durable memory following live-attenuated influenza intranasal vaccination of humans

**DOI:** 10.1101/2025.07.14.664794

**Authors:** Hannah D. Stacey, Lucas Garin-Ortega, Paul G. Lopez, Parham Ramezani-Rad, Sydney I. Ramirez, Farhoud Faraji, Disha Bhavsar, Gina Levi, Florian Krammer, Shane Crotty

## Abstract

Seasonal influenza vaccines are most frequently delivered as intramuscular inactivated vaccines which elicit systemic responses against the immunodominant hemagglutinin (HA) head domain. An intranasally administered, live-attenuated influenza vaccine designed to stimulate mucosal immunity, FluMist, is the sole intranasal vaccine approved in the United States. However, FluMist has lower systemic immunogenicity and efficacy in adults compared to intramuscular formulations. In this study, human mucosal and systemic immunity were examined following seasonal intramuscular or intranasal vaccination. Nasopharyngeal swabs of adenoid tissue were used to longitudinally sample the upper airway. Notably, FluMist induced substantial increases in upper respiratory tract IgG^+^ and IgA^+^ HA-specific memory B cells, which displayed an activated CD27^+^CD21^-^ phenotype. H1, H3, and influenza B virus HA-specific memory B cells were all detected in the upper airway after intranasal immunization and remained elevated at 6-months post-vaccination. Recently activated upper airway memory B cells were not readily detected in intramuscular vaccinees, despite marked elevation of systemic antibody and memory B cells. Thus, despite minimal immune response detected in circulation, live-attenuated influenza vaccine can generate substantial local antigen-specific memory B cell responses in adults. These findings have implications for improving influenza vaccines and for mucosal vaccination against other respiratory pathogens.

**One Sentence Summary:** Longitudinal nasopharyngeal sampling reveals local influenza-specific B cell responses following intranasal but not intramuscular vaccination.

## INTRODUCTION

Influenza viruses pose significant threats to global public health, causing repeated seasonal epidemics that result in millions of infections annually (*1*). In addition, the pandemic potential of influenza viruses from animal reservoirs underscores the need for both improved seasonal vaccines and development of next-generation vaccines with increased breadth (*2*). Inactivated influenza vaccines (IIVs) are administered intramuscularly and contain antigens from two influenza A viruses (IAVs), H1N1 and H3N2, and one or two circulating influenza B virus (IBV) lineages, B/Victoria/2/87-like and B/Yamagata/16/88-like. Current seasonal vaccines generate antibodies against the head domain of the viral entry protein hemagglutinin (HA) (*3*). The induction of antibodies with hemagglutination inhibition (HAI) activity has been demonstrated to correlate with intramuscular influenza vaccine protection (*4*). However, the HA head domain is subject to antigenic drift that renders immunity from prior vaccinations ineffective, necessitating annual reformulation of influenza vaccines.

Enhancing mucosal immune memory or local antibodies at the site of respiratory viral infection represents a potential strategy for improvement of influenza virus vaccines (*5, 6*). The magnitude and kinetics of circulating immunological memory induced by systemic vaccinations for respiratory viral infections of humans have been well established, while local respiratory tract immune memory remains to be well elucidated (*7*). The identification of tissue resident memory B cells (B_Mem_) in the human lung and upper airway after influenza virus, respiratory syncytial virus, or severe acute respiratory syndrome coronavirus 2 (SARS-CoV-2) infection suggests the potential importance of B_Mem_ cells in providing protection from respiratory pathogens (*8–13*). An intranasally delivered live-attenuated influenza vaccine (LAIV), FluMist, designed to replicate in human upper airway has been approved for use in the United States (U.S) since 2003 and in Europe since 2013 (as Fluenz) (*14, 15*). Notably, FluMist remains the only intranasal vaccine approved for use in the U.S. LAIV has repeatedly demonstrated enhanced efficacy compared to IIV in pediatric cohorts but inferior immunogenicity and efficacy in adults (*16–20*). In children, FluMist stimulates systemic HAI antibodies, which are generally not observed in adults (*18, 21*). Additionally, local immunoglobulin A (IgA), tonsillar follicular T helper cell (T_FH_) and local B_Mem_ responses have been observed in pediatric cohorts (*21–23*). A clear understanding of the immune responses elicited by FluMist in adults has remained elusive, in part due to challenges in sampling human mucosal tissues. The capacity of current IIVs to stimulate mucosal immunity also remains incompletely understood (*7, 24*).

As mucosal vaccination strategies become increasingly explored for multiple pathogens, tools for sampling the cellular mucosal response will be essential in evaluating vaccine responses (*25–28*). Newer nasopharyngeal (NP) swab sampling approaches can yield sufficient immune cells to detect B_Mem_, plasma cells (PCs), and T cells (*8, 29*). This minimally invasive sampling technique has the additional key advantage of enabling longitudinal assessment of mucosal immune memory in humans (*8*). Here, we combined conventional and novel approaches to assess local and systemic immunity longitudinally after influenza vaccinations.

## RESULTS

### Study design and influenza vaccine cohorts

To evaluate upper airway and circulating responses in participants who received intramuscular or intranasal influenza vaccination, we obtained matched NP swab, synthetic absorptive matrix (SAM, nasosorption) strip, and peripheral blood (serum, plasma and peripheral blood mononuclear cells (PBMCs)) samples. Samples were collected pre-vaccination, and at 7-, 14-, 30-, 90- and 180-days post-vaccination (Fig. 1A). Participants received either Afluria (intramuscular IIV) or FluMist (intranasal LAIV) containing the 2023-2024 Northern hemisphere strains. Fifty vaccinees were included in the study, with n = 25 per group. Age, sex, and self-reported recent vaccination history were collected (Fig. 1B, S1A-S1C).

**Fig. 1.**
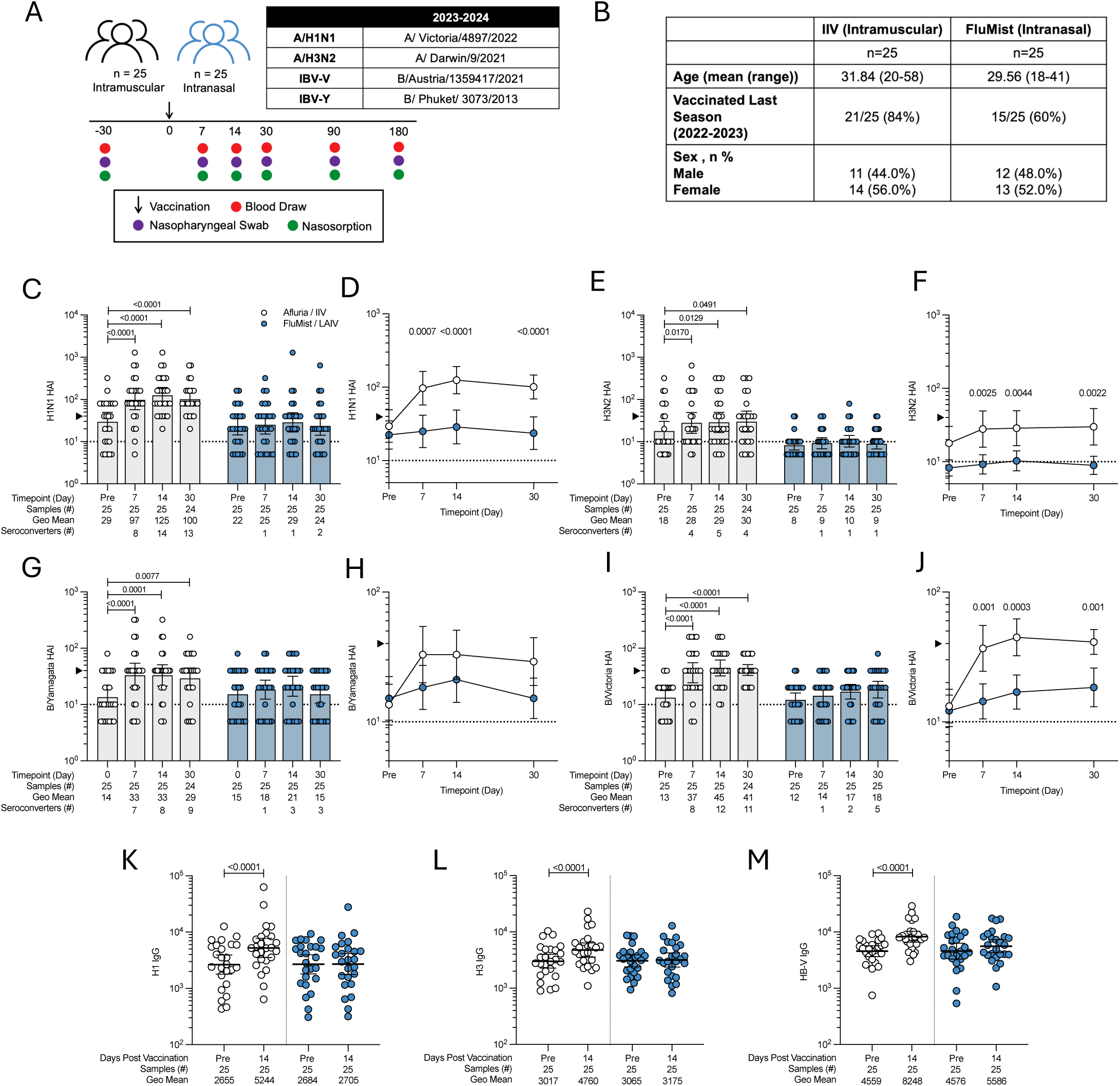
Systemic antibody responses following intramuscular or intranasal influenza vaccination. (A) Study design and sample collection time points. (B) Vaccine cohort characteristics. (C-D) HAI antibody titers against A/Victoria/4897/2022 (H1N1) virus. (E-F) HAI antibody titers against A/ Darwin/9/2021(H3N2) virus.(G-H) HAI antibody titers against B/Phuket/3073/2013 B-Yamagata lineage virus.(I-J) HAI antibody titers against B/Austria/1359417/2021 B-Victoria lineage virus. Comparison of full length (K) H1 HA, (L) H3 HA, and (M) HB-V HA plasma IgG titers pre-vaccination and 14-days post vaccination. IBV-V, influenza B virus Victoria-lineage; IBV-Y, influenza B virus Yamagata-lineage; HB-V, HA of B/Victoria-lineage. (C,E,G,I) Statistical significance of HAI and (K-M) IgG plasma titers relative to baseline were determined using the Wilcoxon matched-pairs signed rank test. (D,F,H,J) Statistical testing between the two vaccines across time points was performed with multiple Mann-Whitney tests with Holm-Šídák’s multiple comparison test. Data are shown as geometric means ± 95% confidence intervals. An HAI titer of 40 is denoted by an arrow. The HAI assay limit of detection (LOD), HAI titer = 10, is shown as a dotted line. Samples with no detectable titer were set to half LOD (HAI titer = 5). LOD for plasma ELISA = 80.

### Systemic antibody responses following IIV or FluMist

Serum A/H1N1 HAI geometric mean titers (GMTs) were significantly elevated following intramuscular IIV vaccination (GMT Pre = 29.4; GMT Day 14 = 124.6; *P* < 0.0001) (Fig. 1C). By 14-days post vaccination, A/H1N1 HAI GMTs increased 4.2-fold (Fig. S2A) and 56% (14/25) of participants had seroconverted (≥4-fold increase in titer). In contrast, A/H1N1 HAI titers were not significantly elevated following FluMist vaccination (Fig. 1C-D). Similar patterns were observed for HAI serum antibody responses to A/H3N2, B/Yamagata/16/88-like (Yamagata), and B/Victoria/2/87-like (Victoria) strains (Fig. 1E-J, S2). HA-binding IgG was also quantified for all subjects and time points. Elevated HA-binding IgG was observed in the IIV cohort (H1, H3, and HB-V (HA of IBV Victoria-lineage), each *P* < 0.0001), whereas HA-binding IgG did not significantly change following FluMist (Fig. 1K-M, Fig. S2E-G). Pre-existing strain-specific antibodies and prior vaccination history impacted serum HAI and plasma IgG responses following IIV administration (Fig. S3, S4). Overall, circulating HAI and anti-HA binding antibodies were selectively increased in subjects receiving intramuscular seasonal influenza vaccination.

### Mucosal HA-binding antibodies following IIV or FluMist vaccination

To evaluate the potential induction of mucosal antibodies following IIV or intranasal FluMist vaccination, we measured antibody titers in the nares, collected by SAM strips pre- and post-vaccination. Consistent with existing literature, IgA was the predominant immunoglobulin (Ig) detected in the nares and was found at a 4:1 to 10:1 ratio to IgG (Fig. 2A-B, S5A-F) (*30, 31*). Nares total IgG increased 2.2-fold in IIV recipients 14-days post-vaccination (Fig. 2A, S5E,S4F).

**Fig. 2.**
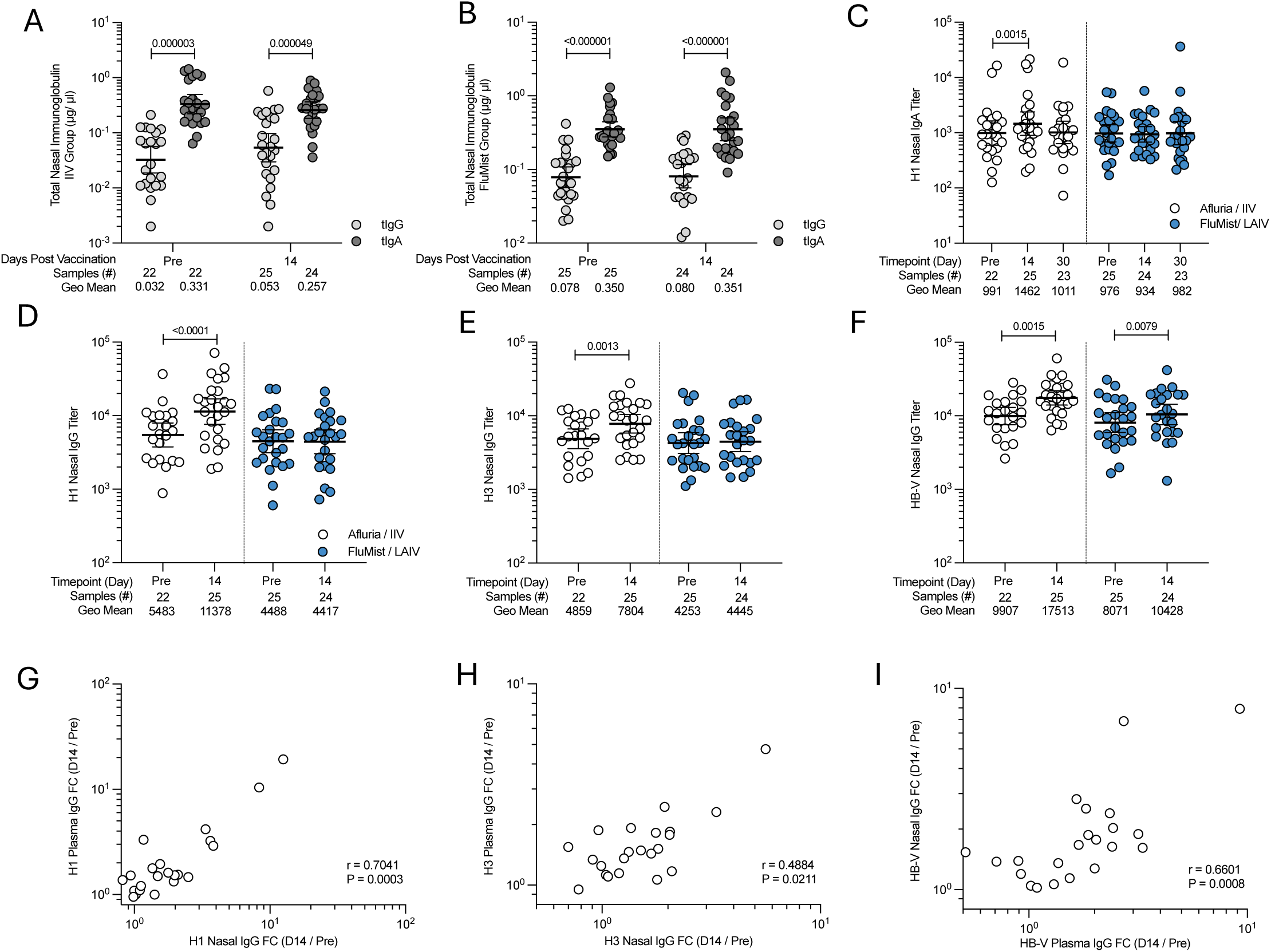
Nasal antibody responses following intramuscular or intranasal influenza vaccination. (A-B) Longitudinal levels of total nasal IgG (light grey) and total nasal IgA (dark grey) from synthetic absorptive matrix samples in the (A) IIV and (B) FluMist cohort. HA-specific binding-IgA and IgG endpoint titers were normalized by total nasal IgA and IgG detected at the matched time point. Wilcoxon matched-pairs signed rank test with Holm-Šídák’s multiple comparison test shown. (C) Comparison of H1 HA nasal IgA adjusted endpoint titers following IIV (white) or LAIV (blue) vaccination. (D-F) Comparison of (D) H1 HA nasal IgG, (E) H3 HA nasal IgG, and (F) HB-V HA nasal IgG adjusted endpoint titers following IIV (white) or LAIV (blue) vaccination. Statistical testing between time points was performed by Wilcoxon matched pairs signed rank test. LOD for antigen-specific nasal immunoglobulin ELISA = 100. All samples are shown but unpaired samples were not considered in the statistical analyses. (G-I) Spearman rank correlations of the day 14 / pre-vaccination fold change in HA-specific plasma IgG compared to the day 14 / pre-vaccination fold change in HA-specific nasal IgG. Spearman correlation r and p values of (G) H1 HA, (H) H3 HA, (I) HB-V HA titers following intramuscular vaccination are shown. (A-F) Data shown as geometric mean ± 95% confidence intervals.

HA-binding IgA and IgG endpoint titers were normalized to total IgA or IgG (µg/ml) in nares samples (adjusted endpoint titer, AET). Increases in mucosal H1 HA-binding IgA were observed in the intramuscular vaccine cohort but not the FluMist cohort (Fig. 2C, S5G,S4H). Increases in mucosal HA-binding IgG for each of H1, H3, and HB-V were observed in IIV recipients, while only an elevation in mucosal HB-V binding IgG was observed following FluMist (Fig. 2D-F). Fold-changes in HA-binding plasma IgG correlated with the increases in nasal HA-binding IgG for each matched HA subtypes in intramuscular vaccinees (H1 r = 0.7041, *P* = 0.0003; H3 r = 0.4884, *P* = 0.0211; HB r = 0.6601, *P* = 0.0008), but not among FluMist recipients (Fig. 2G-I, Fig. S6). Overall, these data indicate FluMist did not induce robust IgG or IgA responses to HA in the nares, while IIV induced a systemic IgG response to all vaccine antigens with a corresponding mucosal IgG elevation consistent with IgG transudation.

### Systemic HA-specific B_Mem_ responses following vaccination

To characterize potential post-vaccination cellular responses, HA-specific B_Mem_ (CD19^+^, IgD^-^, CD27^-/+^, HA^++^) were assessed by flow cytometry using HA probes (Fig. S7). We tested for the presence of HA-specific B cell in the blood and NP swabs of all participants, at all study time points. H1, H3 and HB HA-binding B cells were each measured using two HA probes with separate fluorochromes, for a total of six distinct probes. H1-specific B cells were defined as cell positive for both H1 HA probes (H1 HA^++^), with an identical strategy for H3- and HB-specific B cells (Fig. 3A, Fig. S7). H1, H3, and HB-specific B_Mem_ were detected in the blood of almost all individuals prior to vaccination (Fig. S9A-F).

**Fig. 3.**
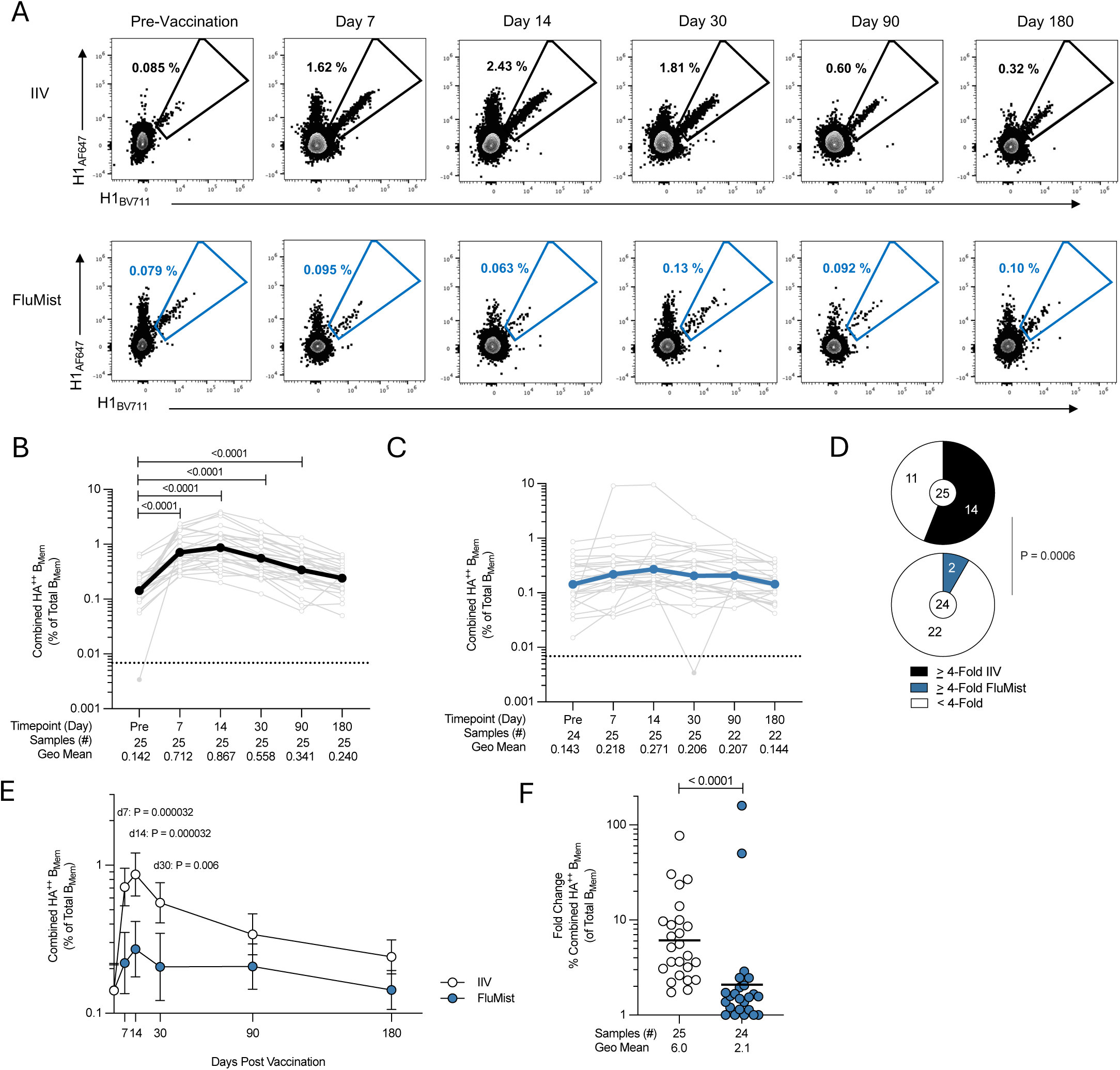
Circulating HA-specific B_Mem_ following seasonal influenza vaccination. (A) Representative gating of longitudinal circulating CD19^+^IgD^-^CD27^+/-^ H1^++^ specific B_Mem_ cells from one representative donor per seasonal vaccine cohort; intramuscular IIV (top) and intranasal FluMist (bottom) vaccination. (B-C) Combined frequency of circulating HA^++^ B_Mem_ following (B) intramuscular or (C) intranasal vaccination. Combined HA^++^ frequency refers to the sum of H1^++^, H3^++^ and HB^++^ frequencies. Light grey joined lines represent individual donors, colored lines represent the geometric mean of the cohort. Dotted lines represent the limit of detection (LOD) determined by the median percentage of 3 antigen-specific B_Mem_ / Total B_Mem_ from all PBMC samples at baseline. The statistical difference between timepoints was assessed by Wilcoxon matched pairs signed rank test. (D) Proportion of IIV (top) or FluMist donors (bottom) with ≥ 4-fold or < 4-fold change in circulating HA^++^ B_Mem._ Fold change was calculated as day 14 frequency/ pre-vaccination frequency. Two-sided Fisher’s exact test shown. (E) Combined frequency of circulating HA^++^ B_Mem_ pre-vaccination and 7-, 14-, 30-, 90- and 180-days post IIV (white) and FluMist (blue) vaccination. Data are displayed as geometric mean ± 95% confidence intervals. Statistical difference between timepoints was assessed by multiple Mann-Whitney test with Holm-Šídák’s multiple comparison test. (F) Day 14/ pre-vaccination fold change of the combined HA^++^ B_Mem_ frequency (% of total B_Mem_) in circulation. The line represents the geometric mean day 14 / pre-vaccination fold change. Fold change ratios < 1 were set to 1. Mann-Whitney test statistic shown.

Combined HA-specific B_Mem_ frequencies (H1+H3+HB B_Mem_) were quantified as a primary metric of vaccine-induced B cell responses. Increased HA-specific B_Mem_ frequencies in blood were detectable 7 days after IIV immunization (*P* < 0.0001, 5.0-fold over pre-vaccination) (Fig. 3B) and peaked 14 days post-vaccination (*P* < 0.0001, 6.1-fold) (Fig. 3B). In contrast, HA-specific B_Mem_ frequencies in blood were not increased > 2-fold at any time point after FluMist (Fig. 3C). The proportion of vaccinees with ≥ 4-fold increases in circulating B_Mem_ frequency was significantly different between the two cohorts: among IIV vaccinees, 56% (14/25) experienced a ≥4-fold increase in circulating HA-specific B_Mem_, while amongst FluMist recipients only 8% (2/24) had a ≥4-fold HA-specific B_Mem_ increase (χ^2^, *P* = 0.0006) (Fig. 3D). The frequency of combined HA-specific B_Mem_ in circulation was significantly higher at 7-, 14-, and 30-days post-IIV compared to FluMist vaccination (Day 7; *P* = 0.000032, Day 14; *P* = 0.000032, and Day 30; *P* = 0.006) (Fig. 3E). The fold change in systemic HA^++^ B_Mem_ following IIV was 6.0 at 14 days post-vaccination (Fig. 3F). Overall, acute and sustained circulating HA-specific B_Mem_ responses were substantial after IIV and not generally observed in FluMist vaccinees.

### Upper airway HA-specific B_Mem_ responses following vaccination

Evaluation of HA-specific B_Mem_ in the upper airway was facilitated by NP swab sampling (Fig. 4A, S8). Before immunization, H1 HA-specific B_Mem_ were detectable in the upper airway of 84% (41/49) of subjects, and 69% (34/49) and 76% (37/49) of participants possessed detectable H3- and HB-specific B_Mem_. 12.3% of HA-specific B_Mem_ in the upper airway were IgA^+^ compared to 1.1% IgA^+^ in blood (pre-immunization medians, Fig. S8).

**Fig. 4.**
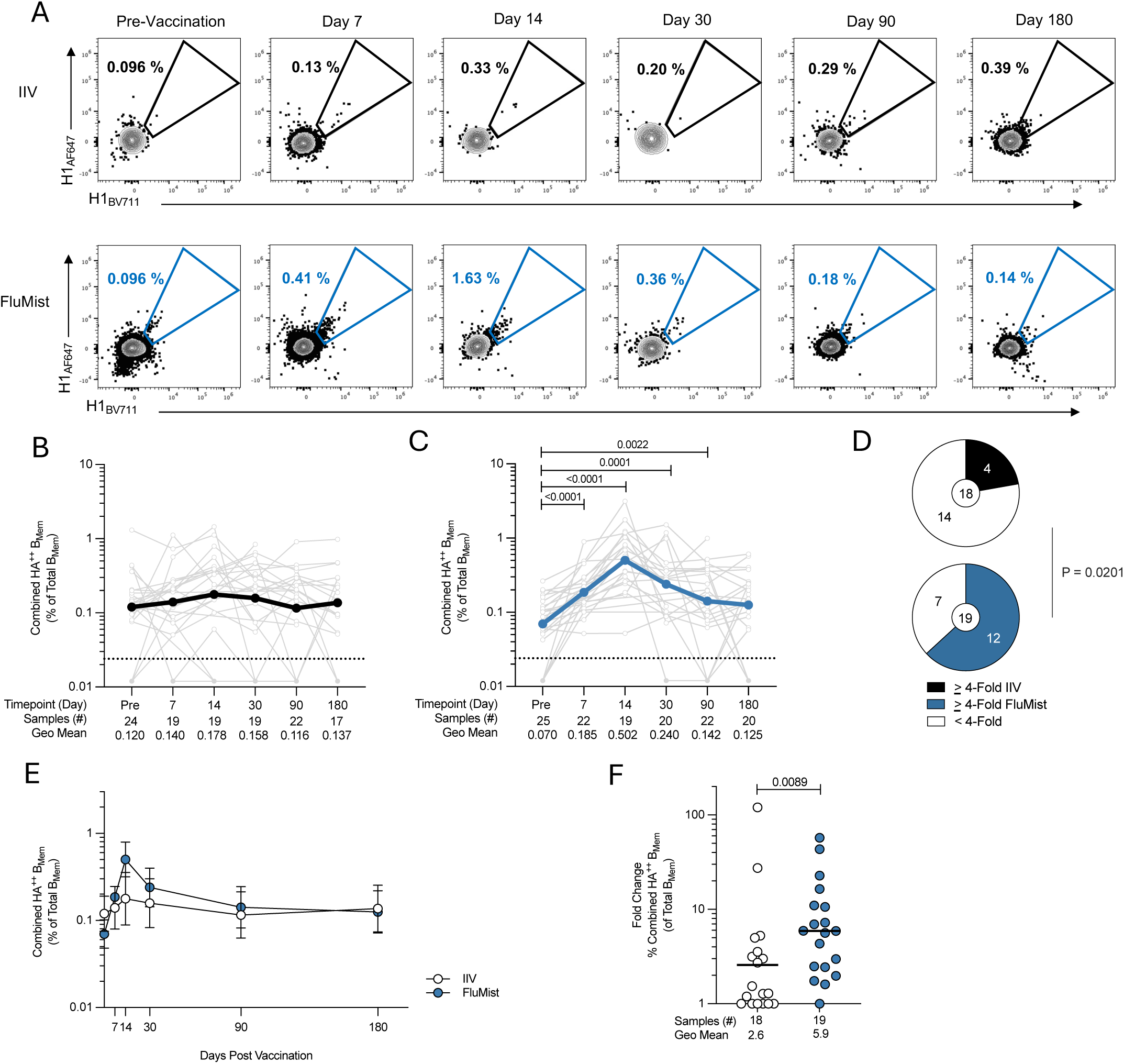
Local HA-specific B_Mem_ in the upper airway following IIV or FluMist vaccination. (A) Representative gating of longitudinal, local CD19^+^CD20^+^ IgD^-^CD27^+/-^ H1^++^ specific upper airway B_Mem_ cells from NP swabs from one representative donor per vaccine cohort. IIV donor (top) and FluMist donor (bottom) vaccination. (B-C) Combined frequencies of upper airway HA^++^ B_Mem_ pre- and post- (B) IIV or (C) FluMist. Combined HA^++^ frequency refers to the sum of H1^++^, H3^++^ and HB^++^ frequencies. Light grey joined lines represent individual donors, colored lines represent the geometric mean of the cohort. Dotted lines represent the limit of detection (LOD) determined by the median percentage of 3 antigen-specific B_Mem_/ Total B_Mem_ from all NP swab samples. The statistical difference between timepoints was assessed by Wilcoxon matched pairs signed rank test. (D) Proportion of IIV donors (top) and FluMist (bottom) donors with ≥ 4-fold or < 4-fold change in URT HA^++^ B_Mem_. Fold change was calculated as day 14 frequency / pre-vaccination frequency. Two-sided Fisher’s exact test shown. (E) Combined frequency of NP HA^++^ B_Mem_ pre-vaccination and 7-, 14-, 30-, 90- and 180-days post IIV (white) and FluMist (blue) vaccination. Data are displayed as geometric mean ± 95% confidence intervals. Statistical difference between timepoints was assessed by multiple Mann-Whitney test with Holm-Šídák’s multiple comparison test. (F) Day 14/ pre-vaccination fold change of the combined HA^++^ B_Mem_ frequency (% of total B_Mem_) in NP swabs. The line represents the geometric mean fold change at 14 days post-vaccination. Fold change ratios < 1 were set to 1. Mann-Whitney test statistic shown.

In participants receiving IIV immunization, geometric mean HA-specific B_Mem_ frequencies in the upper airway did not significantly increase at any time point (Fig. 4B). In contrast, after FluMist immunization, upper airway HA-specific B_Mem_ frequencies were significantly increased (pre-vaccination geometric mean, 0.07%; Day 14, 0.50%) (Fig. 4C). At the peak of the response 2-weeks post vaccination, 18 of 19 FluMist recipients had detectable upper airway specific B_Mem_ responses to individual HA subtypes (Fig. S9). Only 22% (4/18) of IIV participants had ≥4-fold rise in upper airway HA-specific B_Mem_ (Fig. 4D), substantially lower than that observed in circulation after IIV (Fig. 3D). 63% (12/19) of FluMist recipients displayed ≥4-fold rise in HA-specific B_Mem_, significantly more than IIV (χ ^2^, *P* = 0.02).

More granular analysis of HA-specific B_Mem_ by HA subtype after FluMist showed increases in each of H1-, H3- and HB-HA-specific B_Mem_ in the upper airway; H1-binding B_Mem_ increased 3.7-fold after vaccination, H3-binding B_Mem_ increased 3.2-fold, and HB-binding B_Mem_ increased 6.8-fold (Day 14: H1, *P* < 0.0001; H3, *P* = 0.0027; HB, *P* < 0.0001) (Fig. S9). While the absolute frequency of upper airway combined HA-specific B_Mem_ was not significantly different between cohorts (Fig. 4E), day 14 increase fold change was significantly different (χ ^2^, *P* = 0.0089), representing a 5.9-fold change over baseline in the FluMist cohort. (Fig. 4F). In FluMist recipients, upper airway HA-specific B_Mem_ frequencies remained significantly elevated relative to baseline out to 180 days post-vaccination (Day 90, 0.142%, *P* = 0.0001; Day 180, 0.125%, *P* = 0.0022). Upper airway HA-specific B_Mem_ increased in FluMist vaccinees irrespective of receiving an influenza vaccine the previous season (Fig. S10). Overall, these findings indicate that FluMist can induce significant local nasopharyngeal B cell memory, including to each influenza virus strain in the vaccine, and those B_Mem_ responses were largely independent of circulating immune memory.

### Additional T- and B-cell populations in the upper airway

A variety of lymphoid tissue B cell and germinal center (GC) cell populations are readily detectable by NP swabs, including plasma cells (PCs, CD19^+^CD20^-^CD38^hi^), GC B cells (B_GC_, CD19^+^CD20^+^CD38^mid^), and GC-T_FH_ (CD4^+^CXCR5^+^PD-1^hi^) (*8*) (Fig. 5A,5B). No increase in total upper airway PCs was observed post-vaccination (Fig. 5C,5D), nor in total B_GC_ cells or total GC-T_FH_ in the NP swabs (Fig. 5E-5H). No significant increases in HA-specific B_GC_ frequencies were observed at the cohort level (Fig. S8, S11). However, there were more HA-specific B_GC_ responders after FluMist than IIV at 14 days post-vaccination, and this held true for HA-specific B_GC_ responses to all three HA subtypes tested (χ ^2^: H1, *P* = 0.046; H3, *P* = 0.046; HB, *P* = 0.008) (Fig. 5I-K). Thus, some HA-specific GC activity is detectable in response to FluMist in adults suggesting that FluMist, but not IIV, may be capable of eliciting HA-specific upper airway GC responses.

**Fig. 5.**
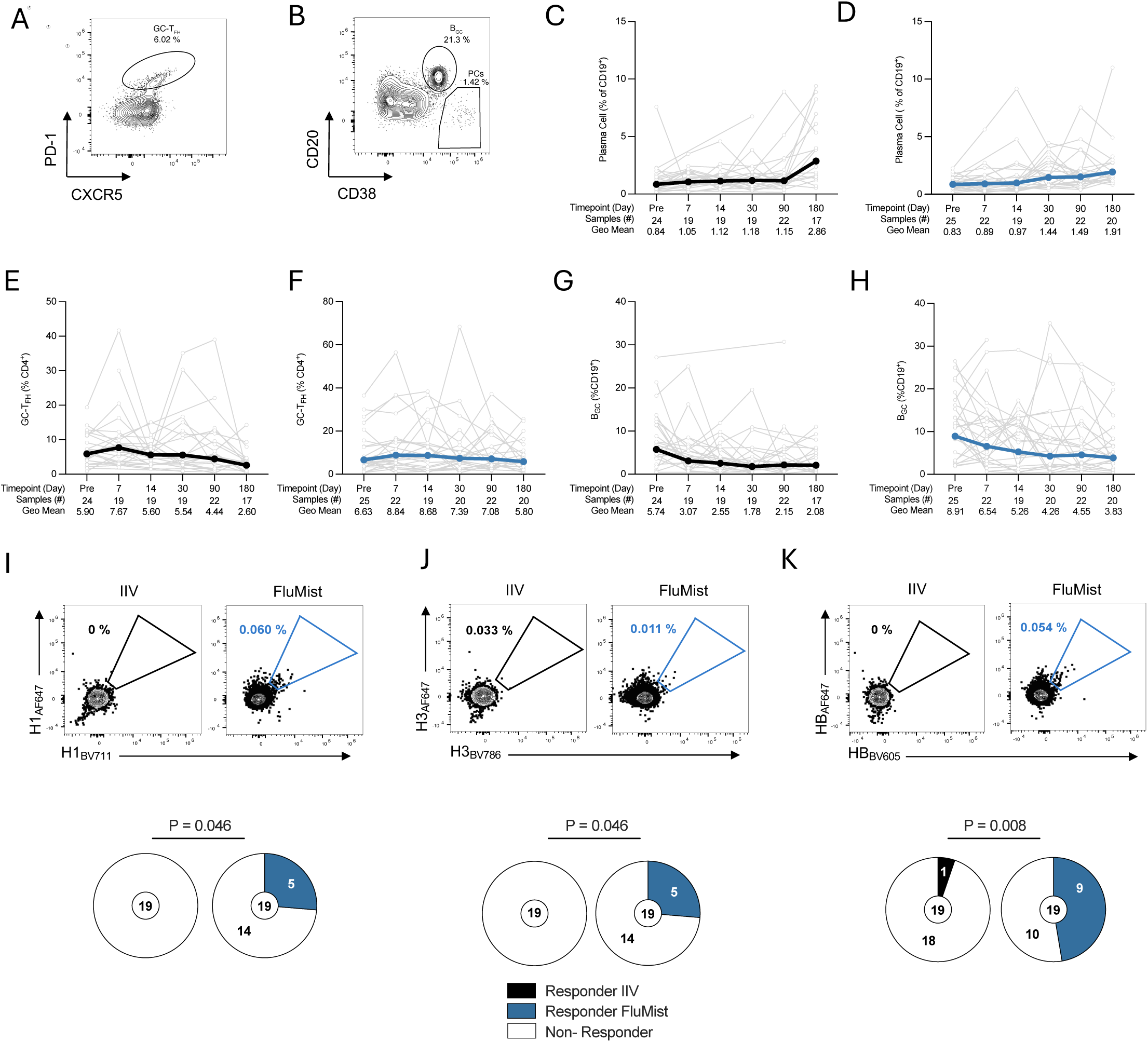
Characterization of other T- and B-cell populations in the upper airway following influenza vaccination. (A) Representative gating of adenoid GC-T_FH_ (CD4^+^CXCR5^+^PD-1^hi^) from NP swabs. (B) Representative gating of B_GC_ and plasma calls from NP swabs. (C-D) Longitudinal data of total plasma cells frequencies (non-antigen specific) in the upper airway following (C) IIV and (D) FluMist. (E-F) Frequency of total GC-T_FH_ in the upper airway following (E) IIV and (F) FluMist. (G-H) Frequency of total adenoid B_GC_ following (G) IIV and (F) FluMist. (I-K) Representative gating of HA-specific B_GC_ and proportion of participants with NP swabs (with ≥ 490 naive B cells) with detectable (I) H1-, (J) H3-, (K) HB-specific adenoid B_GC_ at 14 days post-vaccination. Blue, FluMist recipients with ≥ 3 HA-specific B_GC_. Grey, IIV recipients with ≥ 3-HA-specific B_GC_. (C-H) All light grey lines represent values for individual participants over time, colored lines represent geometric mean for the group. (I-K) Only NP swab samples with ≥ 490 naive (IgD^+^) B cells and ≥ 3 HA-specific (HA^++^) B_GC_ cells for the HA-subtype of interest were included in the analysis. Responder differences assessed by two-sided Fisher’s exact test.

### Isotype switched and activated HA-specific B cells in circulation and the upper airway

Following either vaccination, at all time points, the majority of HA-specific cells in circulation were IgG^+^ (Fig. S13). Circulating memory B cells can be phenotypically characterized utilizing expression of CD21 and CD27; most circulating B_Mem_ have a resting CD27^+^CD21^+^ phenotype (Fig. 6A, S12). A transient population of recently activated CD27^+^CD21^-^ B_Mem_ develops following influenza IIV or other intramuscular vaccinations (*32, 33*). In this study, following IIV, systemic H1-, H3- and HB-specific B_Mem_ with a CD27^+^CD21^-^ phenotype increased rapidly by 7 days post-vaccination, peaking at day 14 (Fig. 6A, 6B, S13). Circulating activated B_Mem_ increases tracked with the peak in the frequency of HA-specific B_Mem_ (Fig. 3B). 14-days after IIV vaccination, 36% of circulating HA-specific (H1+H3+HB) were CD27^+^CD21^-^ B_Mem_, a 6.7-fold increase from baseline (median 5.1%) (Fig. 6B). In contrast, there was no appreciable increase in circulating CD27^+^CD21^-^ B_Mem_ following FluMist vaccination (Fig. S14).

**Fig. 6.**
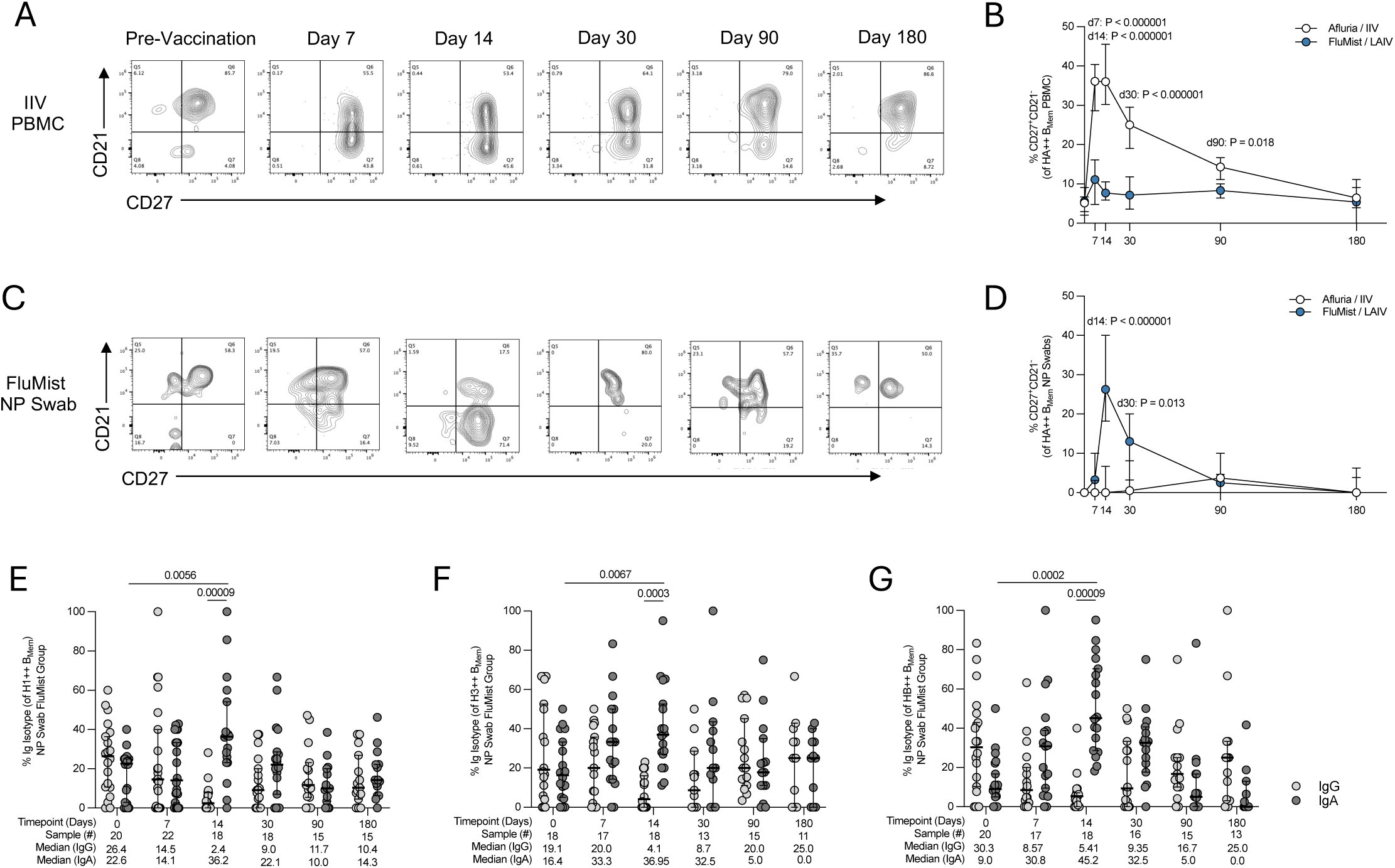
CD27^+^CD21^-^ and immunoglobulin isotype profiling of HA^+^ B_Mem_ elicited by IIV and FluMist. (A) Representative gating strategy for longitudinal CD27/CD21 profiling of circulating HA^++^ B_Mem_ (H1^++^ data displayed) in one donor following IIV administration. (B) Frequency of circulating CD27^+^CD21^-^ HA^++^ cells following IIV (white) or FluMist (blue). (C) Representative gating for longitudinal CD27/CD21 profiling HA^++^ B_Mem_ in the upper airway (H1^++^ data displayed) in one donor following FluMist administration.(D) Frequency of CD27^+^CD21^-^ HA^++^ B_Mem_ in NP swabs following IIV (white) or FluMist (blue). (E-G) Percentage of IgA^+^ (dark grey) and IgG^+^ (light grey) (E) H1, (F) H3, and (G) HB specific upper airway B_Mem_ in FluMist recipients. (B,D, E-G) Data are displayed as median ± 95% confidence intervals. The statistical difference between IgA and IgG HA B_Mem_ at each timepoint was assessed by multiple Mann-Whitney test with Holm-Šídák’s multiple comparison test. The difference within isotypes across timepoints was assessed by Wilcoxon matched pairs signed rank test.

B_Mem_ from the upper airway were next phenotyped. Total activated CD27^+^CD21^-^ B_Mem_ were significantly elevated following FluMist vaccination (Pre to Day 14, *P* = 0.0003) (Fig. 6C, Fig. S12). Among HA-specific B_Mem_, a significant increase in activated CD27^+^CD21^-^ HA B_Mem_ was observed, but only in the FluMist cohort (FluMist vs IIV: H1, *P* = 0.03; H3, *P* = 0.02; HB, *P* = 0.00007) (Fig. 6D, S13-14). Within the FluMist cohort, the median CD27^+^CD21^-^ % at day 14 for all HA subtypes was significantly increased compared to pre-vaccination levels (H1, 0% to 16%, *P* < 0.0001; H3, 0% to 21.3% *P* = 0.005; HB, 0% to 40.6%, *P* < 0.0001).

IgA^+^ HA-specific B_Mem_ increased 14 days post-FluMist in the upper airway, for each of H1, H3, and HB HA-specific B_Mem_ (Pre-vaccination vs Day 14 H1^+^ B_Mem_ *P* = 0.0056; H3^+^ B_Mem_ *P* = 0.0067; HB^+^ B_Mem_ *P* = 0.0002) (Fig. 6E-G). In contrast, proportions of IgA^+^ and IgG^+^ HA-specific B_Mem_ in the upper airway remained unchanged after IIV vaccination (Fig. S13). Given that combined HA^++^ B_Mem_ remained elevated 3.4-fold in the upper airway of FluMist recipients 30 days post-vaccination, the overall data indicate expansions of both IgG^+^ and IgA^+^ B_Mem_ cells in upper airways. Collectively, the increase in recently activated CD27^+^CD21^-^ HA-specific B_Mem_ and the increase in IgA^+^ HA-specific B_Mem_ in the upper airway indicate the selective generation of a local B cell response following FluMist.

## DISCUSSION

Our understanding of mucosal B cell responses following FluMist administration was limited, particularly in adults. In this study we show that by conventional readouts of influenza vaccine immunogenicity—HAI, plasma IgG, B_Mem_ in peripheral blood—FluMist is non-immunogenic in adults. However, within the upper airways, FluMist induced a robust increase of influenza-specific activated B_Mem_ cells, with independent and consistent responses to H1, H3, and HB FluMist antigens, and preferentially of IgA isotype. In contrast, in the periphery increases in HA-specific B_Mem_ and antibodies were selectively detected after IIV. Interestingly, significant anti-HA IgG in the nares was observed only in the IIV cohort, whereas, FluMist induced durable memory in the upper airways out to 6-months post-immunization.

These findings demonstrate that, in a majority of adult recipients, expansion of antigen specific B_Mem_ can be detected in the upper airways 2-weeks after FluMist vaccination. Those responses were undetectable through blood sampling alone, highlighting the importance of relevant tissue sampling for local immune responses. In addition, responses were generated against all vaccine strain HA subtypes, suggesting that FluMist vaccine strains sufficiently replicate to induce local B cell responses in adults. Following FluMist, the elevated activation signature and increase in IgA^+^ isotype HA-specific B_Mem_ cells indicates a likely local tissue origin of these influenza B_Mem_ responses. Additionally, the absence of these responses in the upper airway of intramuscularly administered IIV participants suggests that intranasal delivery and local presentation of antigen is required to elicit or recall mucosal B cell responses.

In addition to HA-specific B_Mem_ responses, upper airway germinal center responses were also observed in the nasopharynx, albeit with more modest magnitudes. Prior work from our group has demonstrated that antigen specific B_GC_ can be detected in NP swabs following SARS-CoV-2 breakthrough infection (*8*). Cold-adaptation of FluMist master donor viruses has attenuated viral replication in the human upper airway, therefore the vaccine response may not be analogous to that observed following natural infection (*34*). The dichotomy between B_Mem_ and B_GC_ responses may also suggest that FluMist is able to induce local B_Mem_ recall responses independently of germinal centers. Alternatively, the adenoids may not be the dominant site of FluMist GCs and GC responses may be occurring in the cervical lymph nodes that drain the nasal cavity mucosa (*35, 36*). The development of less attenuated LAIVs and other novel intranasal influenza vaccine platforms (e.g. subunit adjuvanted) are currently being pursued and may induce more potent GC responses (*37, 38*).

Despite HA-specific B_Mem_ induction in the upper airways, little to no HA-specific antibody was detected following FluMist administration. The uncoupled B_Mem_ and antibody responses observed is consistent with epidemiological data indicating moderate same-year protective efficacy of FluMist in adult cohorts (*17, 19*). Further studies comparing the magnitude, longevity and specificity/breadth of HA-specific B_Mem_ in the upper airways of adult and pediatric donors following FluMist would be beneficial in understanding their contribution to vaccine-mediated protection. A study which examined HA-specific B_Mem_ responses in *ex vivo* tonsils and adenoids by enzyme-linked immunospot (ELISpot) assay demonstrated that pandemic H1N1 infection induced local cross-reactive B_Mem_ in the upper airway (*39*). Similarly, FluMist may prime HA-specific B_Mem_ in the upper airways that can be restimulated to produce antibodies upon re-exposure.

A correlation between systemic IgG and nasal IgG responses in the IIV cohort was observed. Previous studies have demonstrated that HA-specific IgG in nasal wash samples are derived from transudation from plasma rather than local production (*40, 41*). Similar findings were observed in studies examining SARS-CoV-2 hybrid immunity and subsequent intramuscular vaccination (*42, 43*). However, the durability and extent to which transudated IgG in the nasal cavity following intramuscular vaccination contributes to protection provided by seasonal influenza vaccines is not known.

Limitations of this study include the lack of measurement of T cells or cytokines in nasal secretions following vaccine administration, which may have provided additional insights (*44*). Additionally, participants were enrolled in a single influenza season (2023–2024).

Mucosal vaccine strategies are becoming increasingly explored as candidates for multiple pathogens as well as delivery platforms for universal influenza vaccines (*45–47*). These needs highlight the importance of further characterization and improved understanding of immune responses in mucosal tissues. The generation and detection of upper airway B cell responses following intranasal FluMist administration in adults in this study indicates induction and maintenance of B_Mem_ at the mucosa can be an important component of future mucosal vaccines for influenza virus and other respiratory pathogens.

## MATERIALS AND METHODS

### Study population

All samples were collected under a protocol approved by the Institutional Review Board (IRB) of the La Jolla Institute for Immunology (#VD-271). All blood, nasosorption and NP swab samples were collected between September 2023 and May 2024. Healthy adults between 18-65 who had not yet received a 2023-2024 seasonal influenza vaccination were eligible to participate in the intramuscular vaccine cohort. Healthy adults aged 18-49 were eligible to participate in the FluMist cohort as FluMist is not approved for use in individuals over 49 years of age. Participants who received intramuscular vaccination received the 2023-2024 Northern hemisphere Afluria quadrivalent vaccine (CSL Seqirus). Participants who received intranasal vaccination received the 2023-2024 Northern hemisphere FluMist vaccine (AstraZeneca). All participants provided written informed consent for participation in the study.

### Sample collection and processing

All samples were collected by the LJI Clinical Core using standard protocols. Swabs (nasopharyngeal and mid-turbinate) and blood were processed on the same day as collection. For isolation of serum, whole blood was collected into serum separator tubes (SST)(BD vacutainer tubes catalogue no. 367988). SST tubes were rested upright for 15-30 minutes before being spun at 1000 rcf for 10 minutes. Serum was then removed from the upper portion of the tube, aliquoted and stored at -80°C until use. The remainder of the whole blood was collected in ethylenediaminetetraacetic acid (EDTA) tubes (BD vacutainer tubes catalogue no. 366643)(50ml blood draws) or sterile collection bags containing EDTA (100ml blood draws) and kept at room temperature until processing. To separate the plasma, EDTA tubes were spun at 460 rcf (1400 rpm, Rotana-460R, 5699R Rotor) for 7 minutes and whole blood in bags were transferred to 50ml conical tubes and spun at 803 rcf (1850 rpm) for 7 minutes with the break off. Plasma was aliquoted and stored in the -80°C or -30°C until use. For PBMC isolation, 15ml of Histopaque 1077 (Sigma-Aldrich catalogue no.10771) was dispensed into the bottom of a 50ml SepMate tube (StemCell catalogue no. 85450). After plasma was removed from the sample, the remaining blood was diluted to a final volume of 25 ml / 50 ml tube with phosphate buffered saline (PBS) containing 2% heat inactivated fetal bovine serum (FBS). 25ml of blood were gently pipetted into a SepMate tube. Samples were spun at 1200 rcf for 15 minutes at room temperature. Each SepMate tube was inverted, and the contents collected, into a fresh 50ml conical and washed with PBS + 2% FBS. Cells were then spun at 500 rcf for 8 minutes at room temperature. To remove red blood cell contamination, 3-5 ml of ammonium-chloride-potassium buffer (ACK) (Gibco catalogue no. A10492-01) was added to each 50 ml conical tube and allowed to incubate for 5 minutes before being diluted with 10 ml of PBS+ 2% FBS and spun at 450 rcf for 7 minutes. Cell counts and viability check were performed on a Cytek Guava Muse Cell Analyzer (Cytek Biosciences). Five million fresh PBMCs were aliquoted and analyzed by flow-cytometry on the day of collection, the remainder of the cells were cryopreserved in 10% dimethyl sulfoxide (DMSO) in FBS and stored in liquid nitrogen for future use.

Nasopharyngeal swab samples were collected as previously described (*8*). In brief, a flocked swab (Puritan catalogue no. 25-3306-H) was gently inserted into one nostril along the floor of the nasal cavity, parallel to the palette until the posterior NP was reached (∼8 cm). The swab was rotated ∼5-10 times while gently maintaining pressure posteriorly to maintain the swab top within the adenoid pad. Mid-turbinate swab samples were collected by inserting the swab into the nasal cavity in the same fashion as NP swabs, but only ∼3-4 cm into the nasal cavity. The inferior turbinate mucosa was then sampled by rotating the swab in a circular motion (∼5-10 full rotations) while maintaining pressure against the inferior turbinate. Swab samples were collected bilaterally using the same swab to sample the adenoid/ NP from both the left and right nares, or to sample the left and right inferior turbinates. Swabs were placed in a 15 ml falcon tube containing 2ml of Roswell Park Memorial Institute (RPMI) media with 10% heat inactivated FBS, 5% Penicillin-Streptomycin (Gibco catalogue no. 15140-122), 5% GlutaMax (Gibco catalogue no. 35050-61) and stored at 4°C until processing. Following collection, swabs were vortexed to release cells from the swab. To further release cells, swabs were placed on a pre-moistened 40µm filter placed over an open 50 ml conical tube and were rinsed with an additional 10ml of supplemented RPMI media (10% FBS, 5% GlutaMax, 5% Pen-Strep). Single cell swab suspensions were stored at 4°C until staining.

Nasosorption samples were collected by having the participants place a synthetic absorptive matrix (SAM) strip (Hunt Developments catalogue no. NSFL-FX1-11) into the left or right naris and applying external pressure on the same nostril to hold the strip in place for 1-2 minutes. Nasosorption samples were stored at -80°C until processing. To process SAM strips, samples were thawed on ice for 20 minutes. The SAM strip was cut from the handle and placed into a labelled 1.7 ml Eppendorf tube. The SAM strip was saturated with 400 µl of cold antibody stabilization buffer (BD catalogue no. 559148), incubated with eluent for at least 1 minute and vortexed for 1 minute to dissociate sample from the SAM. The strip was then transferred to a Costar Spin-X centrifuge tube insert (Costar catalogue no. 9301) in a new 2 ml tube and spun at 452 rcf (Eppendorf 5427R Centrifuge) for 10 minutes at 4 °C. The eluted nasosorption sample was split into two aliquots and stored at -80 °C until use.

### Viruses

Influenza A (A/Victoria/4897/2022(H1N1) and A/Darwin/9/2021(H3N2)) and B (B/Phuket/3073/2013 and B/Austria/1359417/2021) viruses were propagated in 10-12 day old embryonated chicken eggs. Influenza A viruses were incubated for 48 hours at 37 °C. Influenza B viruses were incubated for 72 hours at 33 °C Allantoic fluid was harvested and titrated using turkey or guinea pig red blood cells prior to use.

### Hemagglutination inhibition (HAI) assay

20µL starting volume of serum was treated with 0.5×(starting volume) of 8mg/mL tosyl phenylalanyl chloromethyl ketone (TPCK)-treated trypsin (Sigma-Aldrich) and incubated at 56°C for 30 minutes. After cooling to room temperature, 3x(starting volume) of 11mM potassium periodate (Sigma-Aldrich) was added and incubated for 15 minutes at room temperature. 3x(starting volume) of 1% glycerol-PBS was then added and incubated for 15 minutes. Lastly, 2.5x(starting volume) of 0.85% PBS was added to all samples. Trypsin-heat-periodate inactivation was utilized to reduce the chance of false positives. Serum was diluted in a series of 2-fold serial dilutions in 96-well V-bottom plates (Sarstedt catalogue no. 82.1583.001). An equal volume of influenza virus, adjusted to 8 hemagglutination units (HAU)/50µL diluted in 1x PBS, was added to each well of the plate. Sera and virus were incubated for 30 minutes at room temperature. After this incubation, 50µL of 0.5% turkey red blood cells (Lampire Biologicals) or 0.75% guinea pig red blood cells (GPRBCs) in 1x PBS (Lampire Biologicals) were added to all wells and incubated at 4°C for 45 minutes. GPRBCs were used for H3N2 A/Darwin/9/2021 due to lack of viral activity observed with turkey red blood cells. After incubation with red blood cells the plates were tilted to observe the hemagglutination inhibition. The HAI titer was determined by taking the reciprocal dilution of the last well that contained non-agglutinated red blood cells. For GPRBC plates, the HAI titer was determined as the last dilution with a “halo” of the same size as PBS and virus only wells. Samples with no detectable activity were assigned to half the limit of detection (HAI=5).

To confirm assay consistency, normal control goat serum (FR-1377) and positive control influenza Reference Goat Antiserum (FR-1863-A/Victoria/4897/2022(H1N1), FR-1827-A/Darwin/9/2021(H3N2), FR-1828-B/Austria/1359417/2021, and FR-1829-B/Phuket/3073/2013) were obtained through the International Reagent Resource, Influenza Division, WHO Collaborating Center for Surveillance, Epidemiology and Control of Influenza, Centers for Disease Control and Prevention, Atlanta, GA, USA. In addition, serum for the same participant at the various timepoints were run on the same plate for each virus, together with the negative and positive control serum.

### HA IgG Plasma Enzyme-Linked Immunosorbent Assay (ELISA)

Recombinant HA protein was obtained from AcroBiosystems (H1 HA catalogue no. HA1-V52H8; H3 HA catalogue no.HA2-V52H6; HB HA catalogue no. HAE-V52H3). Corning 96-well half area plates (Corning catalogue no.3690) were coated with 1 ug/ml of HA in PBS overnight at 4°C. The following day, coating buffer was removed and plates were blocked with 3% skim milk in PBS containing 0.05% Tween-20 (PBS-T) for 1.5 hours at room temperature (100µl / well). Plasma samples were diluted in 1% skim milk in 0.05% PBS-T beginning at a starting dilution of 1:80, followed by 11, 2-fold serial dilutions. Samples were incubated for 1.5 hours at room temperature. Plasma samples were run in technical duplicates for each time point on the same plate and a positive control of pooled serum from vaccinated donors was used on each plate to ensure consistency across assay days. Following the incubation, plates were washed five times with PBS containing 0.1% Tween-20 (0.1% PBS-T) (100µl /well). To detect IgG, the secondary IgG-horseradish peroxidase (HRP) antibody (Hybridoma Reagent Laboratory catalogue no. HP6043-HRP) was diluted 1:16,000 in 1% skim milk in 0.05% PBS-T. After the final wash, 50 µl per well of secondary antibody was added to the plate and allowed to incubate for 1.5 hours at room temperature. Plates were washed five times with 0.1% PBS-T before adding tetramethylbenzidine (TMB) substrate (ThermoFisher Scientific catalogue no. 34021) (30µl /well) and allowing plates to incubate for 5 minutes. The reaction was stopped with an equivalent volume of 2N sulfuric acid. Plates were read at 450nm on a Spectramax M2E plate reader with the RevB adapter (Molecular Devices). Plate backgrounds were determined by calculating the average of 12 blank wells on the plate. Endpoint titers for each sample were determined using background subtracted OD (optical density) values. Asymmetric sigmoidal 5PL interpolation (Graphpad Prism) was then used to determine the dilution at which the background-corrected OD reached 0.1 OD.

### Nasosorption (synthetic absorptive matrix) Total IgG/ IgA ELISA

To account for potential sampling/ antibody concentration variation during SAM collection, antigen specific HA-binding nasal IgA and IgG endpoint titers were normalized to total IgA or IgG (µg/ml) (adjusted endpoint titer, AET).NUNC Maxisorp ELISA plates (ThermoFisher catalogue no. 442404) were coated overnight at 4°C with capture IgG (Hybridoma Reagent Labs catalogue no.HP6046P) or capture IgA antibodies (Hybridoma Reagent Labs catalogue no. HP6111P) at a 1:200 dilution (10ug/ml) in PBS (100µl /well). The following day the coating buffer was removed, and plates were blocked for 1.5 hours at room temperature with 100µl/ well of PBS containing 10% FBS and 0.2% Tween-20. Nasosorption samples were diluted in the blocking buffer at a starting dilution of 1:100, followed by 11, 2-fold dilutions across the plate. Polyclonal human immunoglobulins were used as standards for determination of total IgG and IgA. For IgG, a polyclonal standard (Jackson ImmunoResearch Labs, #009-000-003) beginning at 0.7µg/ml was diluted 2-fold across the plate. For IgA, ChromPure Human Serum IgA (Jackson ImmunoResearch Labs, #009-000-011) beginning at 22 µg/ml was used as a standard. Samples were allowed to incubate for 1.5 hours at room temperature. Plates were then washed 3 times with PBS containing 0.05% Tween-20 (0.05% PBS-T) (100µl /well). Secondary antibodies were diluted in 10% FBS and 0.2% Tween-20. For IgG detection an IgG-HRP secondary (Hybridoma Reagent Laboratory, #HP6043-HRP) was used at 1:16,000.For IgA, IgA-HRP secondary (Hybridoma Reagent Laboratory, #HP6123-HRP) was used at 1:8000. Secondary antibody was incubated for 1.5 hours at room temperature before plates were washed again 3 times with 0.05% PBS-T. 100µl/ well of TMB substrate (ThermoFisher Scientific catalogue no. 34021) was incubated for 5 minutes and the reaction was stopped by adding an equivalent volume of 2N sulfuric acid. The unadjusted OD values of the standard were used to interpolate the total IgG or IgA concentrations using asymmetric sigmoidal 5PL interpolation (Graphpad). For each sample the OD value that fell in the middle of the linear range of the dilution curve was selected for interpolation. The interpolated concentration was multiplied by the selected dilution factor and expressed in µg/ml.

### Nasosorption (synthetic absorptive matrix) Antigen Specific ELISA

Corning 96-well half area plates (Corning #3690) were coated with 1 µg/ml of HA in PBS overnight at 4°C (30µl/well). The following day, coating buffer was removed. For IgG detection plates were blocked for 1.5 hours at room temperature with 3% Milk in 0.05% PBS-T. For IgA detection 3% milk in PBS was used as a blocking buffer. Nasosorption samples were diluted at 1:40 in 1% milk in 0.05% PBS-T for IgG plates and 3% bovine serum albumin (BSA) in 0.1% PBS-T for IgA detection. Samples were 2-fold diluted 11 times and incubated for 1.5 hours at room temperature. Samples were run in technical duplicates the same plate and a positive control of pooled serum from vaccinated donors was used on each plate to ensure consistency across assay days. After incubation plates were washed five times with 0.1% PBS-T (100µl/well). For IgG detection a goat anti-human IgG-biotin antibody (ThermoFisher catalogue no.A18821) was diluted at 1:32,000 in 1% milk in 0.05% PBS-T (50µl/well). For IgA detection, an anti-human IgA-biotin antibody was diluted at 1:4000 (Southern Biotech catalogue no. 2050-08) in 3% BSA in 0.1% PBS-T (50µl/well). Both IgG and IgA plates were incubated for 1.5 hours at room temperature before washing five times with 0.1% PBS-T. Pierce high-sensitivity streptavidin-HRP (ThermoFisher catalogue no. 21130) was used at 1:32,000 (50µl/well) in the respective diluent buffers as previously stated. The streptavidin was incubated for 1.5 hours and plates were washed five times with 0.1% PBS-T. To develop the plates, 30µl of one step TMB-Ultra (ThermoFisher catalogue no. 34029) was added to each well and allowed to develop for ∼4 minutes. The reaction was stopped with an equivalent volume of 2N sulfuric acid. Plates were read at 450nm on a Spectramax M2E plate reader (Molecular Devices). Plate backgrounds were determined by calculating the average of 12 blank wells on the plate. Endpoint titers for each sample were determined using background subtracted OD values. Asymmetric sigmoidal 5PL interpolation (Graphpad Prism) was then used to determine the dilution at which the background-corrected OD reached 0.1 OD. Adjusted endpoints for antigen specific IgG and IgA from nasosorption samples were calculated by dividing the endpoint by the total IgG or IgA concentration from the corresponding sample.

### Detection of HA-specific B cells by flow cytometry

Detection of influenza HA-specific B cells was performed using B cell probes. Recombinant HA was purchased from AcroBiosystems (H1 HA, HA1-V52H8; H3 HA, HA2-V52H6; HB HA HAE-V52H3). HA was biotinylated using the EZ-Link-NHS-biotinylation kit (ThermoFisher catalogue no. 21935). Recombinant HA was biotinylated with a 25-fold molar excess of biotin. To enhance specificity, HA probes were made using two fluorochrome colors for each protein. H1 HA was incubated either Alexa Fluor 647 (BioLegend, catalogue no. 405237) or BV711 (BD, catalogue no. 563262), H3 was incubated with Alexa Fluor 647 or BV786 (BD, catalogue no. 563858) and influenza B HA (Victoria lineage) was incubated with either Alexa Fluor 647 or BV605 (BD, catalogue no. 563260). All streptavidin-biotinylated HA complexes were made at a 4:1 molar ratio. Probes were prepared individually then combined into a master mix containing brilliant stain buffer (BD Biosciences, catalogue no. 566385)) and 5µM of free D-biotin (Avidity Biosciences, catalogue no. BIO-200).

### Flow cytometry

Samples were pre-stained with LIVE/DEAD Fixable Blue Stain Kit 1:500 in PBS (ThermoFisher Scientific, catalogue no. L34962) and 5 µl / sample of human Fc block (BioLegend, catalogue no. 422302) in PBS at room temperature for 15 minutes. NP swabs were stained with 100µl of antigen probe master mix containing 250 ng/ HA-subtype/ fluorochrome (500 ng / subtype). PBMCs were stained with 50 µl of the same probe cocktail. Samples were stained with probe for 1 hour at 4°C prior to the addition of surface staining antibodies. Samples were incubated with surface antibodies (Table S1) in Brilliant Stain Buffer (BD, catalogue no. 566349) for 30 minutes at 4°C. Data was acquired on a 5-laser Cytek Aurora spectral cytometer (Cytek Biosciences) and analyzed using FlowJo v10 (BD Bioscience).

To address inherent variability in NP swab sample collection, to be included in the analysis, NP swabs had to meet a threshold of ≥490 naïve B (Live, CD45^+^,CD19^+^, CD20^+^,IgD^+^). This threshold was used to delineate if an adequate adenoid sample had been collected. Any sample with less that 490 naïve B cells was not included in any downstream analysis. The limit of detection (LOD) for NP swabs was determined by the median percentage of 3/ Total B_Mem_ from all NP swab samples (LOD, 0.024 %). The limit of detection for PBMC samples was calculated as the median percentage of 3/ Total B_Mem_ from all PBMC samples at baseline (LOD, 0.0069 %). For the combined HA^++^ (H1+H3+HB) analysis, in cases of samples where an individual HA subtypes had less than 3 HA double positive cells, the frequency of that HA subtype was not included in the final combined HA frequency. Samples with less than 3 total HA specific cells were set to the ½ LOD (NP Swab, 0.012%; PBMC, 0.0034%).For individual HA subtype analyses, if a sample had >490 naïve B cells but < 3 individual HA-specific cells, samples at that time point were set to ½ LOD for the respective sample type. Samples with < 3 HA++ cells are depicted as a colored circles on the graph. For inclusion in B_Mem_ isotyping and CD21/CD27 phenotyping analyses, in addition to requiring ≥ 490 naïve B cells, a threshold of 3 HA double positive cells for the subtype of interest was used.

### Statistics

Statistical analysis was performed in GraphPad Prism 10 (GraphPad Software v10.4). All flow cytometry data were analyzed with FlowJo (v10). Additional details can be found in the figures and corresponding figure legends.

## ACKNOWLEDGEMENTS

We would like to thank all the donors who participated and made this work possible. We are extremely grateful to the Clinical Studies Core staff at LJI; Gina Levi, Jasmine Cardenas, Noemi Chavez for sample collection. We thank Isabella Simmons, Ana De Wallau John, and Brittany Nunn for processing peripheral blood samples. We would also like to extend our thanks to Christina Kim and Hayley Simon for assistance with IRB protocol submissions and documentation. In addition, we thank clinical study coordinators, Alex Herrera, Crystal Valdez, and Suzy Aguilar for their support. We also thank all the other LJI core services who provided their knowledge, assistance and guidance in the completion of this study.

## AUTHOR CONTRIBUTIONS

Conceptualization H.D.S., L.G.O., S. I. R., and P.R.R., F.F., and S.C. Clinical activities were supervised, and most samples were collected by G.L. Sample processing and flow cytometry were performed by H.D.S., L.G.O, and P.G.L., L.G.O. also carried out serological assays (ELISA) and data analysis with oversight provided by H.D.S. Essential viral reagents were provided by D.B. and F.K. H.D.S and L.G.O., contributed to figure generation. H.D.S and S.C. wrote the original draft. Supervision and funding were provided by S.C. All authors reviewed and edited this paper.

## DATA AND MATERIALS AVAILABILITY

All data associated with this study are included in the paper or the supplementary materials.

## SUPPLEMENTARY FIGURE LEGENDS

**Fig. S1.**
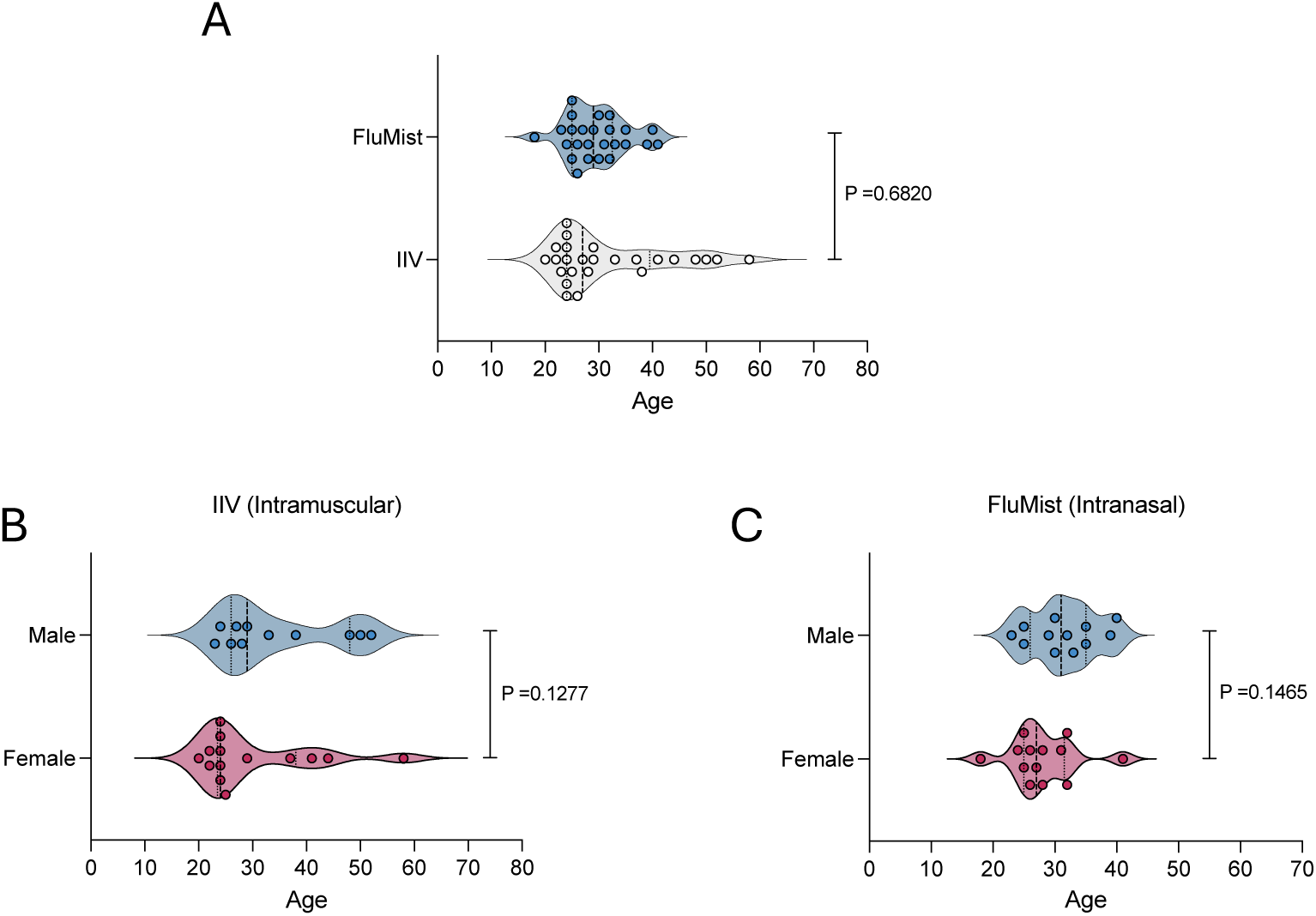
Study participant demographics. **(A)** Comparison of ages across the IIV and FluMist vaccine cohorts. **(B,C)** Comparison of ages between the sexes within each vaccine cohort; **(B)** IIV, **(C)** FluMist. Data is displayed as violin plots showing the median age across vaccine cohorts and sex within vaccine cohorts. Data were analyzed for statistical significance by Mann-Whitney test.

**Fig. S2.**
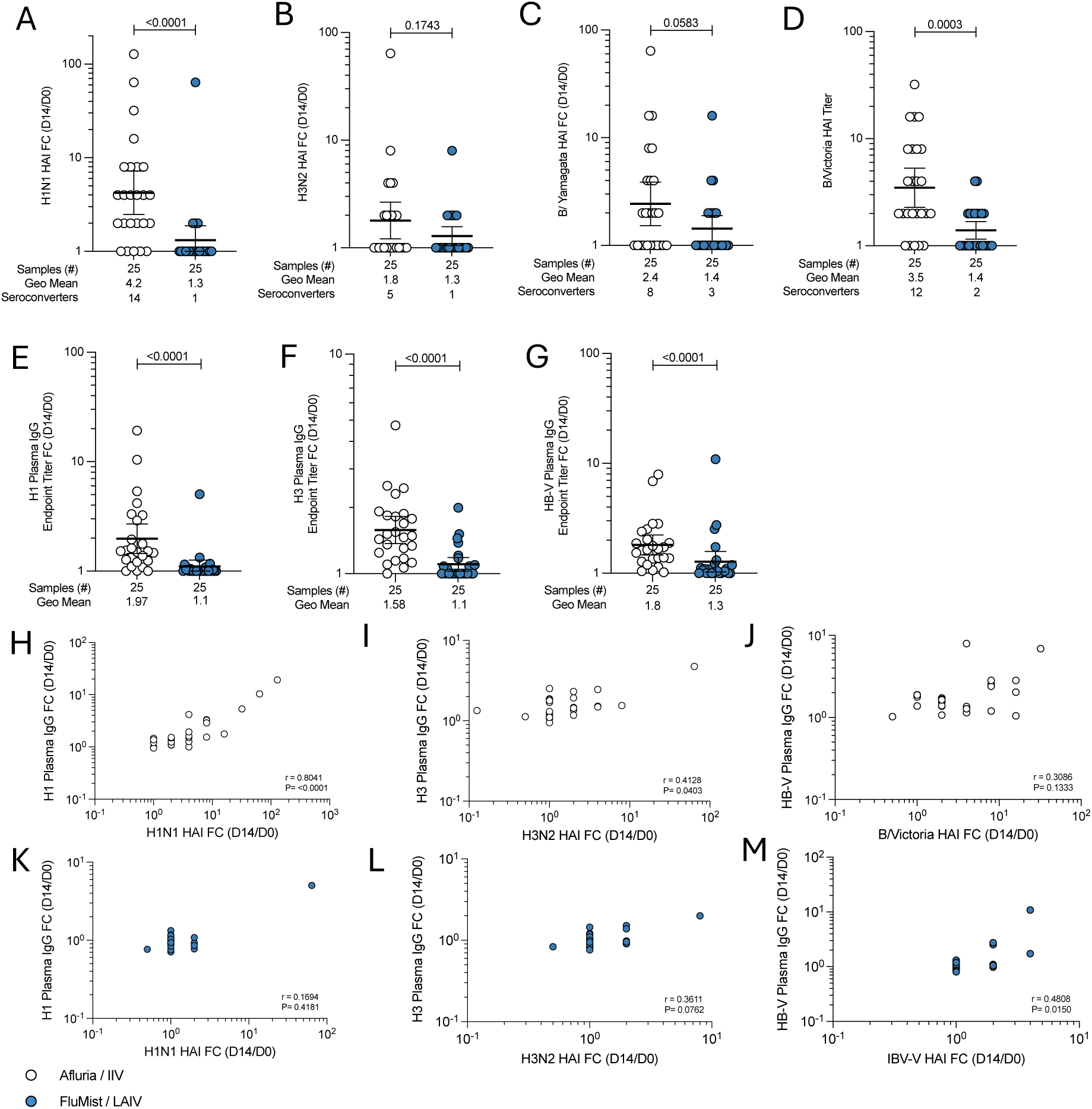
Systemic antibody responses elicited by intramuscular or intranasal vaccination. Fold change of HAI antibody responses between pre-vaccination and 14 days post-vaccination against **(A)** A/H1N1 **(B)** A/H3N2 **(C)** B/Yamagata and **(D)** B/Victoria–lineage virus strains from the 2023-2024 Northern hemisphere vaccine. Fold change of IgG plasma endpoint titers for **(E)** H1 HA **(F)** H3 HA and **(G)** HB-V HA. **(A-G)** Data are displayed as geometric mean ± 95% confidence intervals. **(H-M)** Spearman rank-order correlations of HAI titer fold changes at and IgG plasma titer fold changes from pre-vaccination vs. 14-days post vaccination; **(H)** H1N1 and H1 HA titers, **(I)** H3N2 and H3 HA titers, and **(J)** B/Victoria and HB-V HA titers following IIV; **(K)** H1N1 and H1 HA titers, **(L)** H3N2 and H3 HA titers, and **(M)** B/Victoria and HB-V HA titers following FluMist vaccination. FC, Fold Change; HAI, hemagglutination inhibition; IgG, immunoglobulin IgG; IBV-V, influenza B virus Victoria-lineage, IBV-Y, influenza B virus Yamagata-lineage. Statistical significance of fold change data **(A-G)** was assessed by Mann-Whitney test, correlations were determined by Spearman rank-order correlations **(H-M)**.

**Fig. S3.**
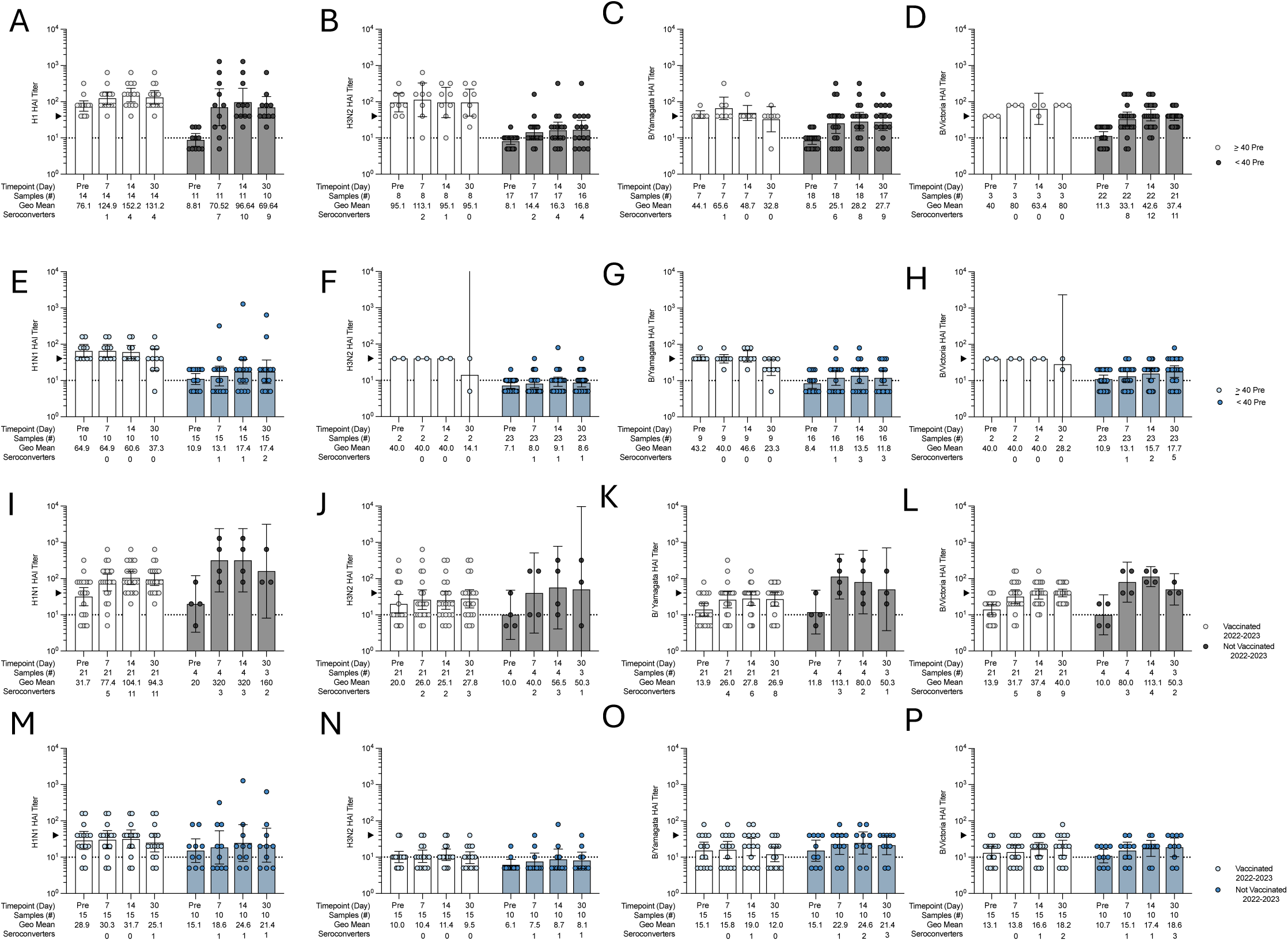
The impact of vaccination history and pre-existing antibody titers on systemic antibody responses. An HAI titer of 40 has been demonstrated to be a seroprotective threshold. The seasonal vaccine cohorts were subdivided into individuals who had an HAI ≥ 40 **(A-H)**. Data was additionally analyzed by splitting the cohorts based on self-reported seasonal influenza vaccine status for the influenza season prior to the study **(I-P)**. HAI titers were measured against the matched **(A,E,I,M)** A/H1N1, **(B,F,J,N)** A/H3N2, **(C,G,K,O)** B/Yamagata-lineage, **(D,H,L,P)** B/Victoria-lineage 2023-2024 Northern Hemisphere vaccine strains. **(A-D)** HAI titers in the IIV cohort based on baseline HAI titer < 40 (grey) or ≥ 40 (white). **(E-H)** HAI titers in the LAIV cohort based on baseline HAI titer < 40 (blue) or ≥ 40 (white). **(I-L)** HAI titers in the IIV cohort based on self-reported vaccination history during the influenza season preceding the study, where vaccinated (white) or not vaccinated (grey) in the previous season. (M-P) HAI titers in the LAIV cohort based on self-reported vaccination history during the influenza season preceding the study, where vaccinated (white) or not vaccinated (blue) in the previous season. IIV participants with pre-vaccination titers < 40 had increased titers and seroconversion rates compared to those with baseline titers > 40 against the vaccine strain. These differences were most pronounced for A/H1N1, B/Yamagata and B/Victoria titers. A/H3N2 responses were 3.3-fold higher 14-days post vaccination in participants with baseline titers <40. B/Victoria seroconversion was only observed in participants with baseline titers <40. Similar trends were observed when IIV cohort HAI responses were assessed based on self-reported vaccination history. In contrast, prior vaccination and pre-existing HAI titers did not significantly impact seroconversion or the magnitude of the HAI response following LAIV administration. Data are displayed as geometric means ± 95% confidence intervals. The seroprotective threshold of HAI = 40 is shown on the y-axis as a black triangle. The HAI limit of detection (LOD = 10) is shown as a dotted black line. Participants with HAI titers below the LOD were set to an HAI titer of 5.

**Fig. S4.**
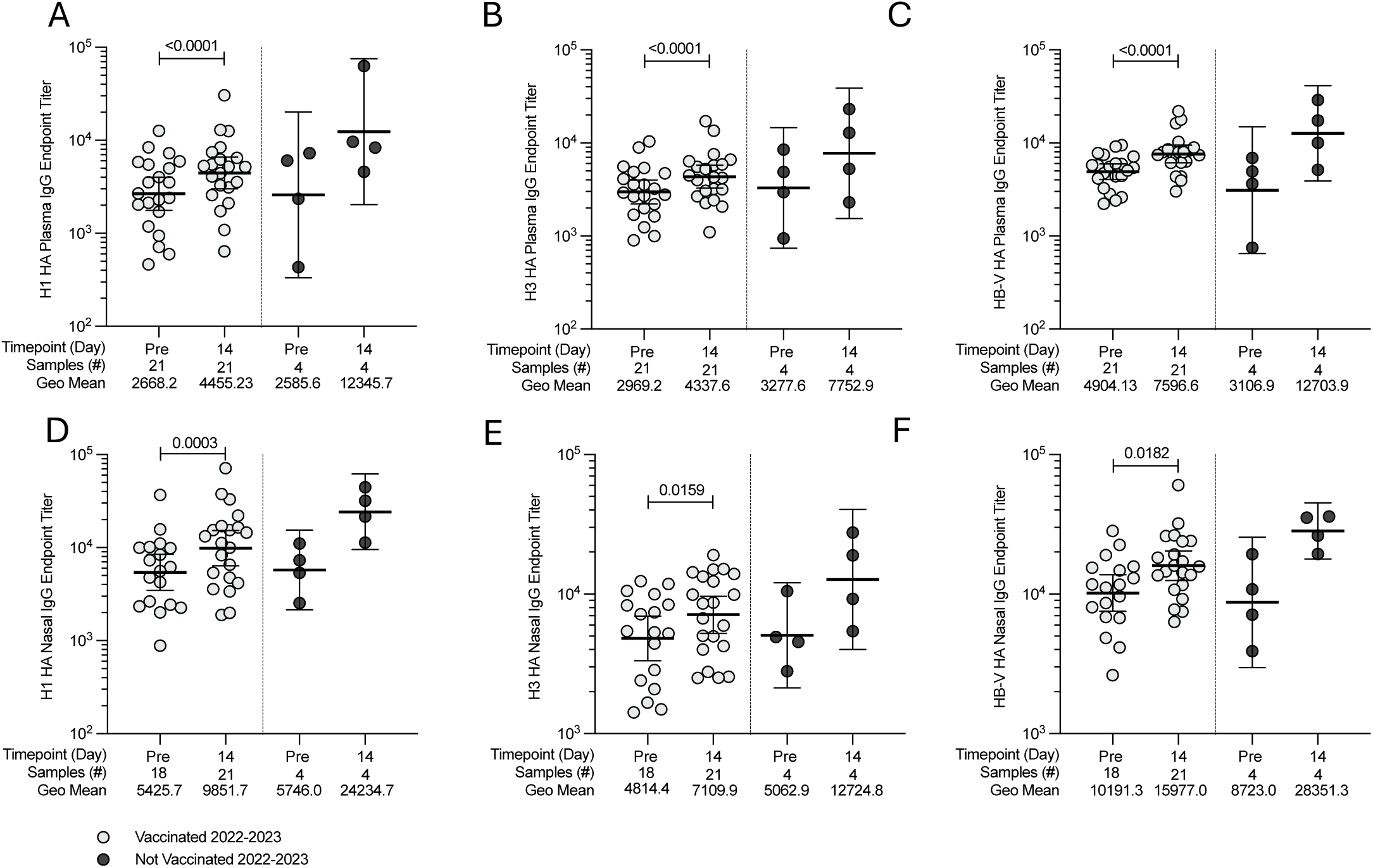
The impact of vaccination history on nasal IgG responses. **(A-C)** Plasma binding IgG **(A)** H1 **(B)** H3 and **(C)** HB IgG endpoint titers in the IIV cohort based on self-reported vaccination history during the influenza season preceding the study, where vaccinated (light grey) or not vaccinated (dark grey) in the previous season. **(D-F)** Nasal **(D)** H1 **(E)** H3 and **(F)** HB IgG adjusted endpoint titers in the IIV cohort based on self-reported vaccination history during the influenza season preceding the study, where vaccinated (light grey) or not vaccinated (dark grey) in the previous season. Statistical significance between the timepoints was assessed by Wilcoxon matched pairs signed rank test. Participants in the intramuscular vaccine cohort who reported not receiving influenza vaccination in the prior season had greater (but non-significant) day 14-fold changes for plasma and nasal HA-binding IgG. All data are displayed as geometric mean ± 95% confidence intervals.

**Fig. S5.**
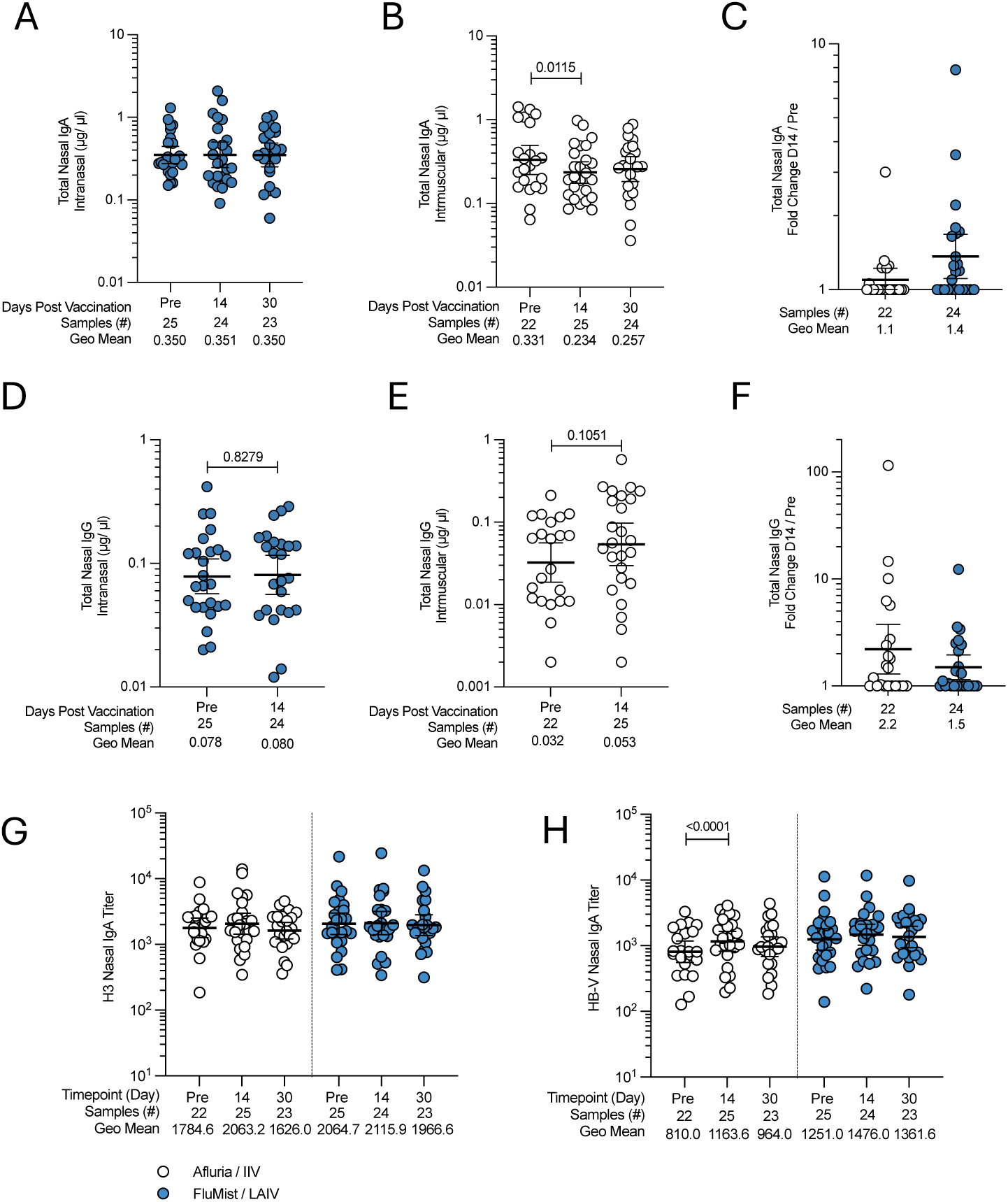
Total nasal immunoglobulin detection collected by synthetic absorptive matrix sampling. **(A)** Total IgA detected pre-vaccination,14- and 30- days post-vaccination for LAIV vaccinees. **(B)** Total IgA detected pre-vaccination, 14- and 30-days post-vaccination for IIV vaccinees. **(C)** Comparison of total nasal IgA fold change from pre-vaccination and 14 days post-vaccination for both vaccine cohorts. **(D)** Total IgG detected pre-vaccination and 14- days post-vaccination for LAIV vaccinees. **(E)** Total IgG detected pre-vaccination and 14-days post-vaccination for IIV vaccinees. **(F)** Comparison of total nasal IgG fold change from pre-vaccination and 14-days post-vaccination for both vaccine cohorts. **(G-H)** Comparison of **(G)** H3 HA nasal IgA and **(H)** HB-V HA nasal IgA adjusted endpoint titers following intramuscular (white) or intranasal (blue) vaccination. **(A,B,D,E)** Statistical significance of total immunoglobulin levels between timepoints was determined by paired Wilcoxon test, all samples are shown but unpaired samples were not considered in the statistical analysis. **(C,F)** Day 14- fold changes in total IgG and IgA were compared by Mann-Whitney test. **(G-H)** Statistical significance between timepoints was determined by Wilcoxon matched pairs signed rank test. All samples are shown but unpaired samples were not considered in the statistical analysis. All data are displayed as geometric mean ± 95% confidence intervals.

**Fig. S6.**
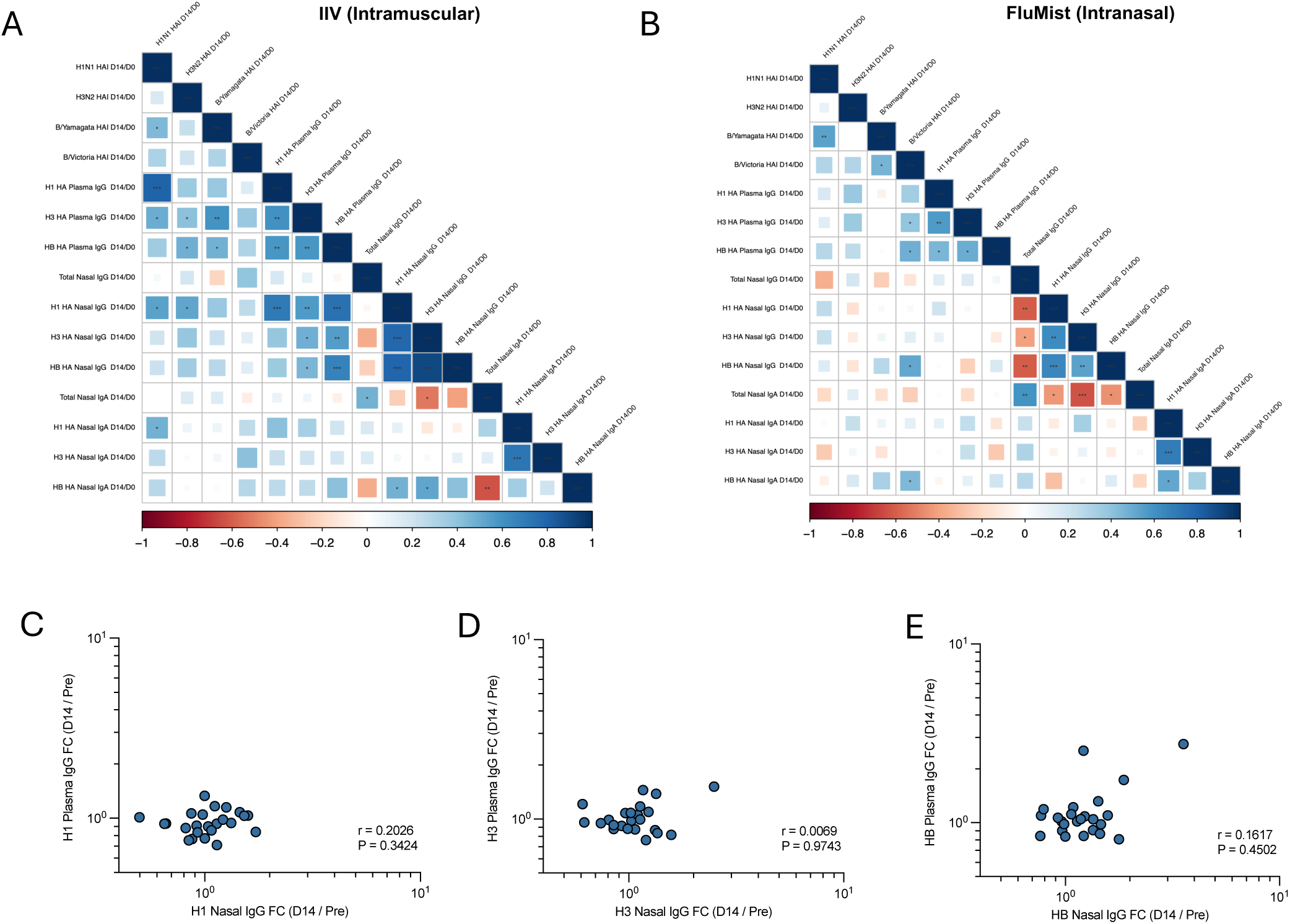
Comparison of upper airway and systemic humoral responses. Spearman rank-order correlation matrix comparing day 14 / pre-vaccination fold changes in nasal and systemic serological data from the **(A)** IIV cohort and **(B)** FluMist cohort. **(C-E)** Spearman rank correlation of the day 14- fold change in HA-specific plasma IgG compared to the fold change in HA-specific nasal IgG. Spearman correlation of **(C)** H1 HA, **(D)** H3 HA, **(E)** HB HA titers following intranasal vaccination. Correlation matrices were made using the Spearman rank order correlation values (r) which are shown as colors ranging from red (-1) to blue (1.0) and square size. P-values are shown as black stars where * p <0.01, ** p<0.01, *** p<0.001.

**Fig. S7.**
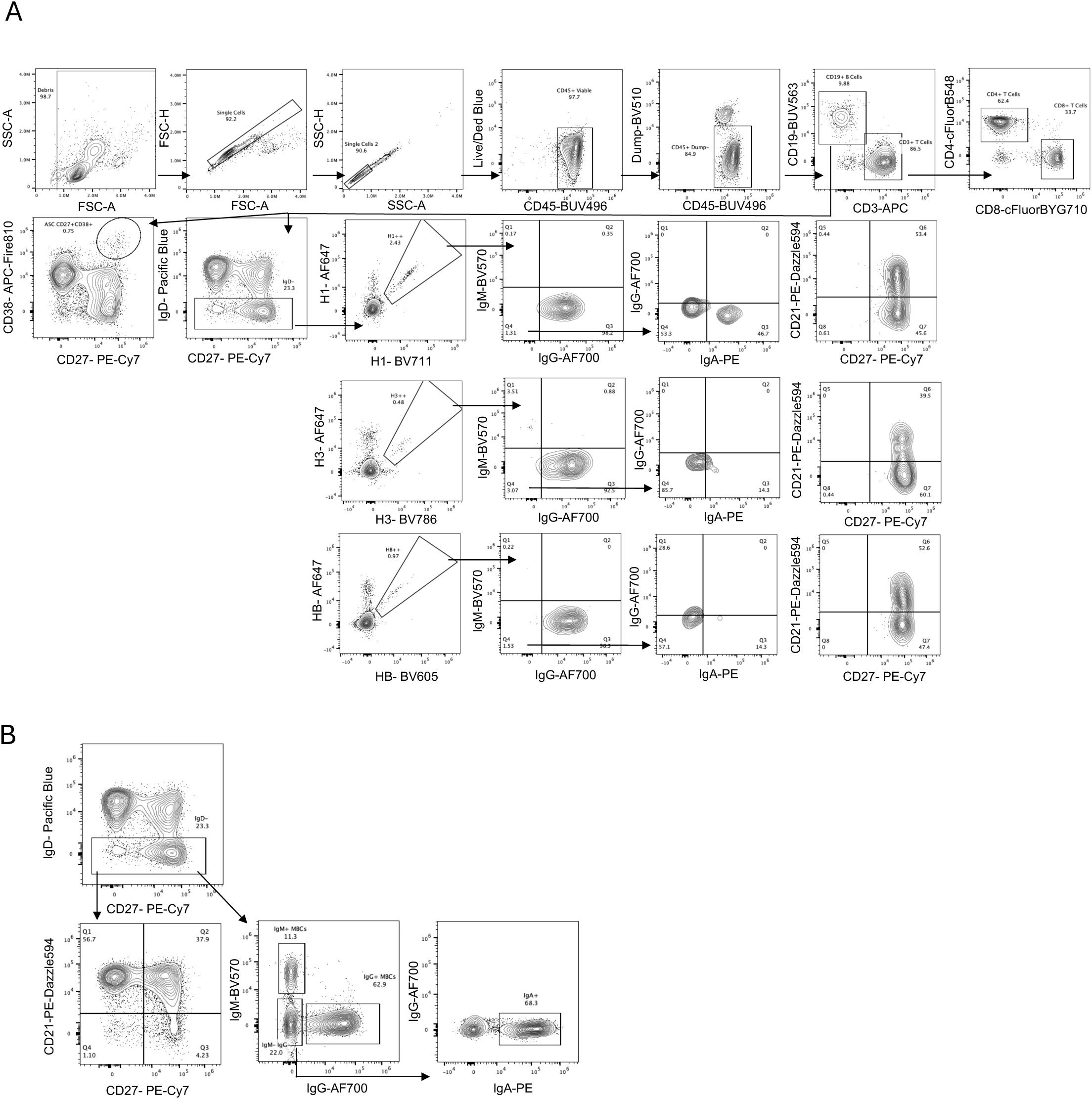
Flow cytometry gating strategies for phenotype and isotype assignment of circulating T cells and BMem from PBMC. Gating strategy for circulating **(A)** HA-specific and **(B)** total B_Mem_ populations from PBMC, including activation status (CD27/CD21) and isotype.

**Fig. S8.**
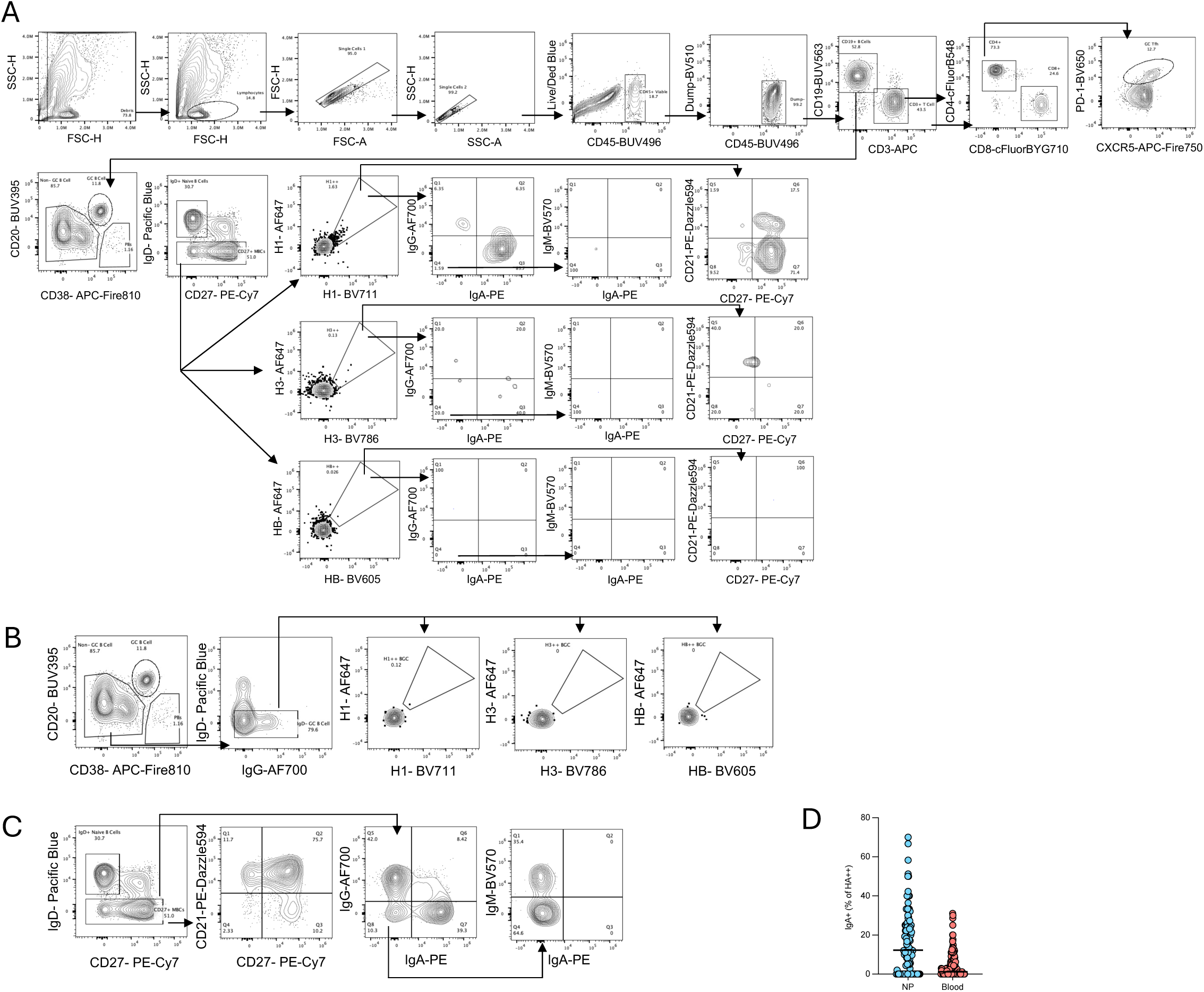
Flow cytometry gating strategies for phenotype and isotype assignment of B and T cells from NP swabs. Gating strategy for **(A)** HA- specific B_Mem_ from NP swabs, **(B)** adenoid HA- specific B_GC_, **(C)** CD21/CD27 phenotyping and isotype assignment of total class switched (IgD^-^) B_Mem_ populations in NP swabs. **(D)** Frequency of pre-vaccination IgA^+^ HA-specific B_Mem_ in NP swabs and peripheral blood agnostic of vaccine cohort; data are displayed as a median for each anatomic site.

**Fig. S9.**
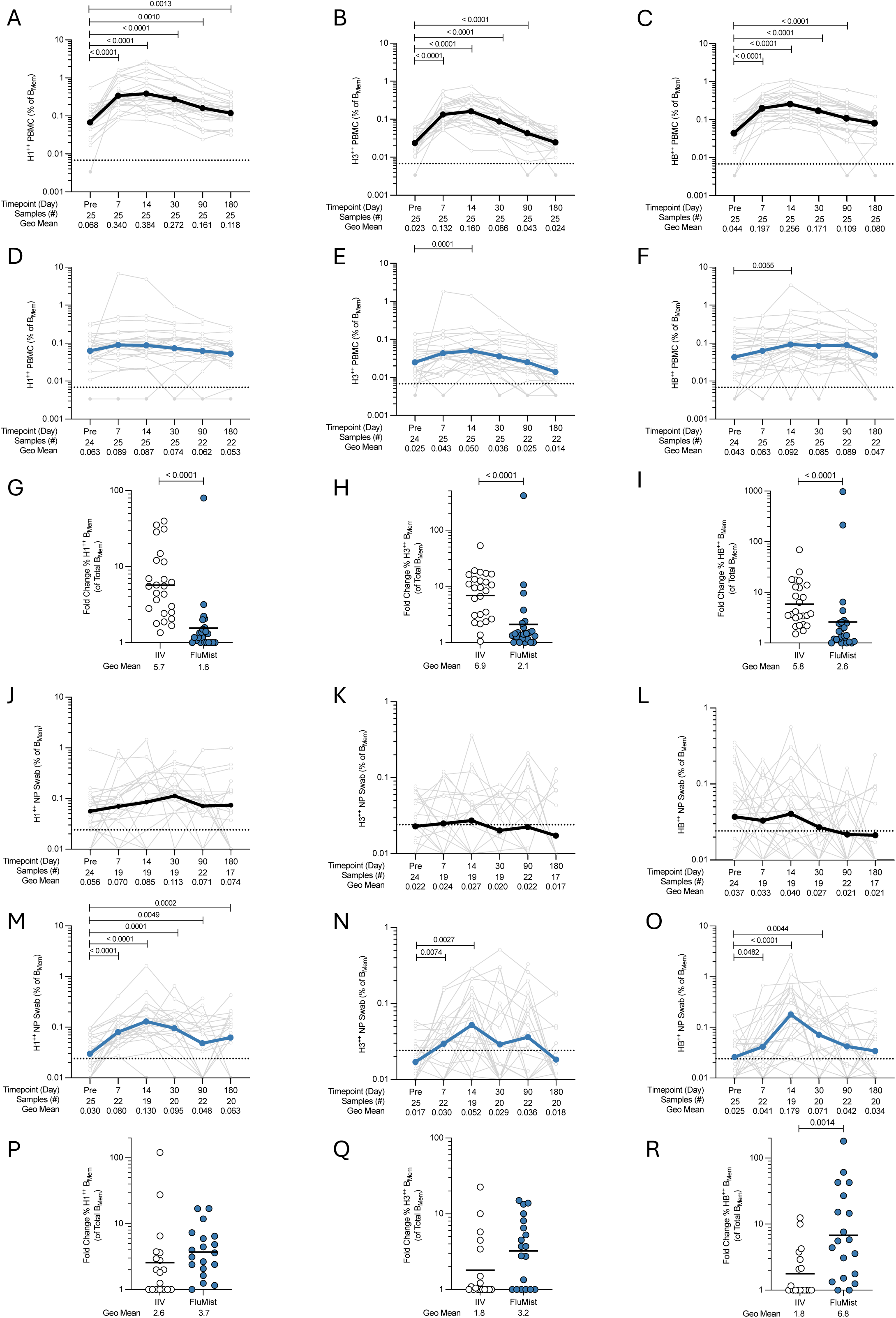
HA-specific B_Mem_ frequencies from PBMC and NP swabs analyzed by HA subtype. **(A-C)** Frequency of circulating **(A)** H1-, **(B)** H3-, and **(C)** HB-specific B_Mem_ in IIV recipients. **(D-F)** Frequency of systemic **(D)** H1-, **(E)** H3-, and **(F)** HB-specific B_Mem_ in FluMist recipients. **(G-I)** Day 14/ pre-vaccination fold change in circulating **(G)** H1-, **(H)** H3-, **(I)** HB-specific B_Mem_ frequencies. Frequency of local **(J)** H1-, **(K)** H3-, and **(L)** HB-specific B_Mem_ in NP swabs of IIV recipients. **(M-O)** Frequency of local **(M)** H1-, **(N)** H3-, and **(O)** HB-specific B_Mem_ in NP swabs of FluMist recipients. **(P-R)** Day 14/ pre-vaccination fold change in circulating **(P)** H1-, **(Q)** H3-, **(R)** HB-specific B_Mem_ frequencies. Light grey joined lines represent values for individual donors, colored lines represent the geometric mean for the cohort. Only successful adenoid swab samples, with naïve B cell (IgD^+^) count of ≥ 490 were included in the analysis. Samples with ≥ 490 naïve B cells but < 3 antigen-specific cells for the HA-subtype of interest were set to the ½ the limit of detection (LOD) and are depicted by colored symbols. Dotted lines represent the limit LOD determined by the median percentage equivalent to 3 antigen-specific B_Mem_/ Total B_Mem_ in all samples at the baseline timepoint (PBMC data) and the median of 3 antigen-specific B_Mem_/ Total B_Mem_ from all NP swab samples. **(A-F, J-O)** Statistical significance between timepoints was determined by Wilcoxon matched pairs signed rank test. **(G-I, P-R)** Statistical differences in the D14/ Pre-Vaccination fold change was determined by two-tailed Mann-Whitney test. Data are displayed as the geometric mean of the fold change. Fold change ratios < 1 were set to 1.

**Fig. S10.**
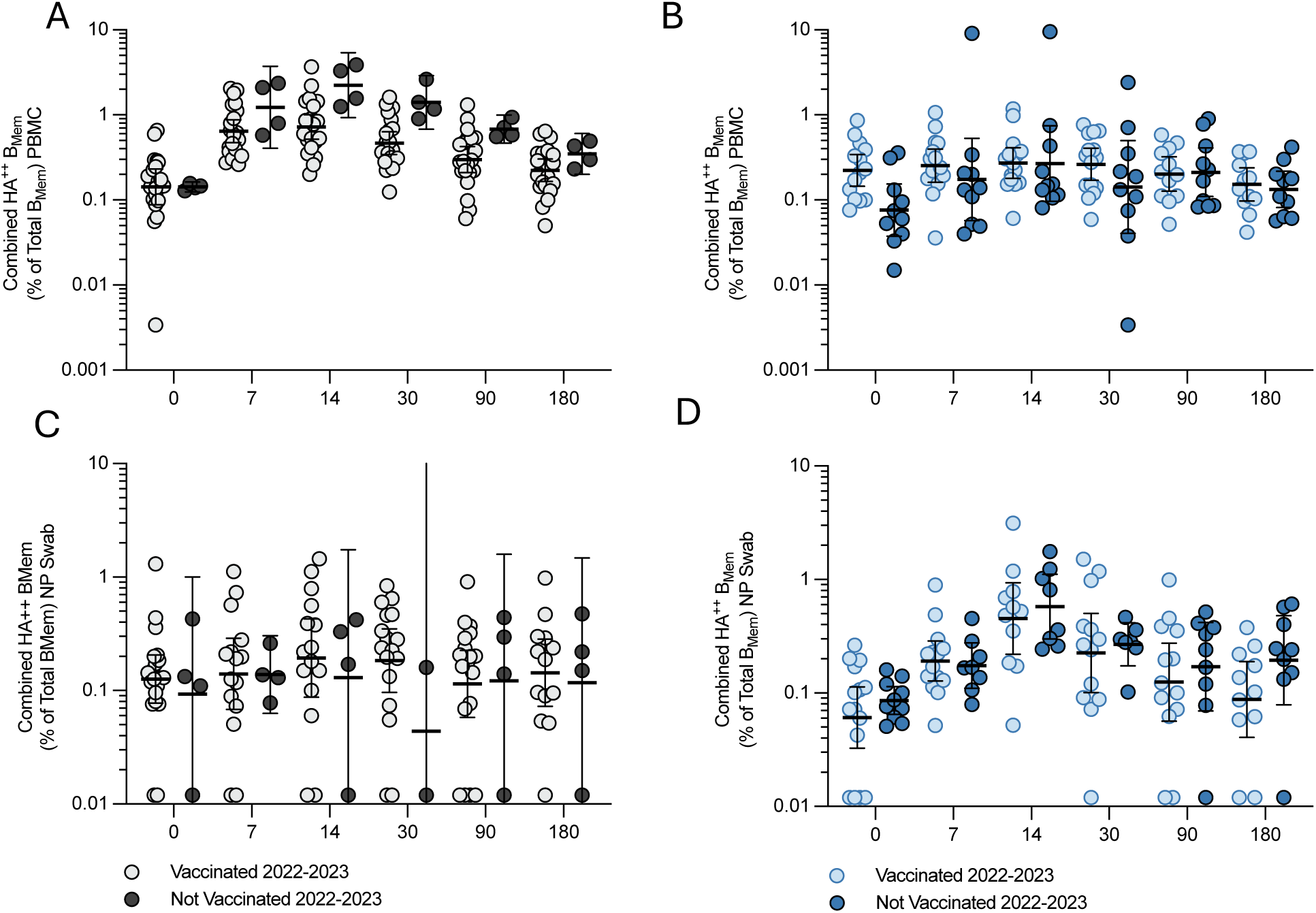
Impact of past season influenza vaccination on NP and circulating HA-specific B_Mem_ induction. Combined HA^++^ B_Mem_ frequency in the peripheral blood following **(A)** IIV and **(B)** FluMist. Combined HA^++^B_Mem_ frequency in the upper airway following **(C)** IIV and **(D)** FluMist. Data are displayed as geometric mean ± 95% confidence intervals.

**Fig. S11.**
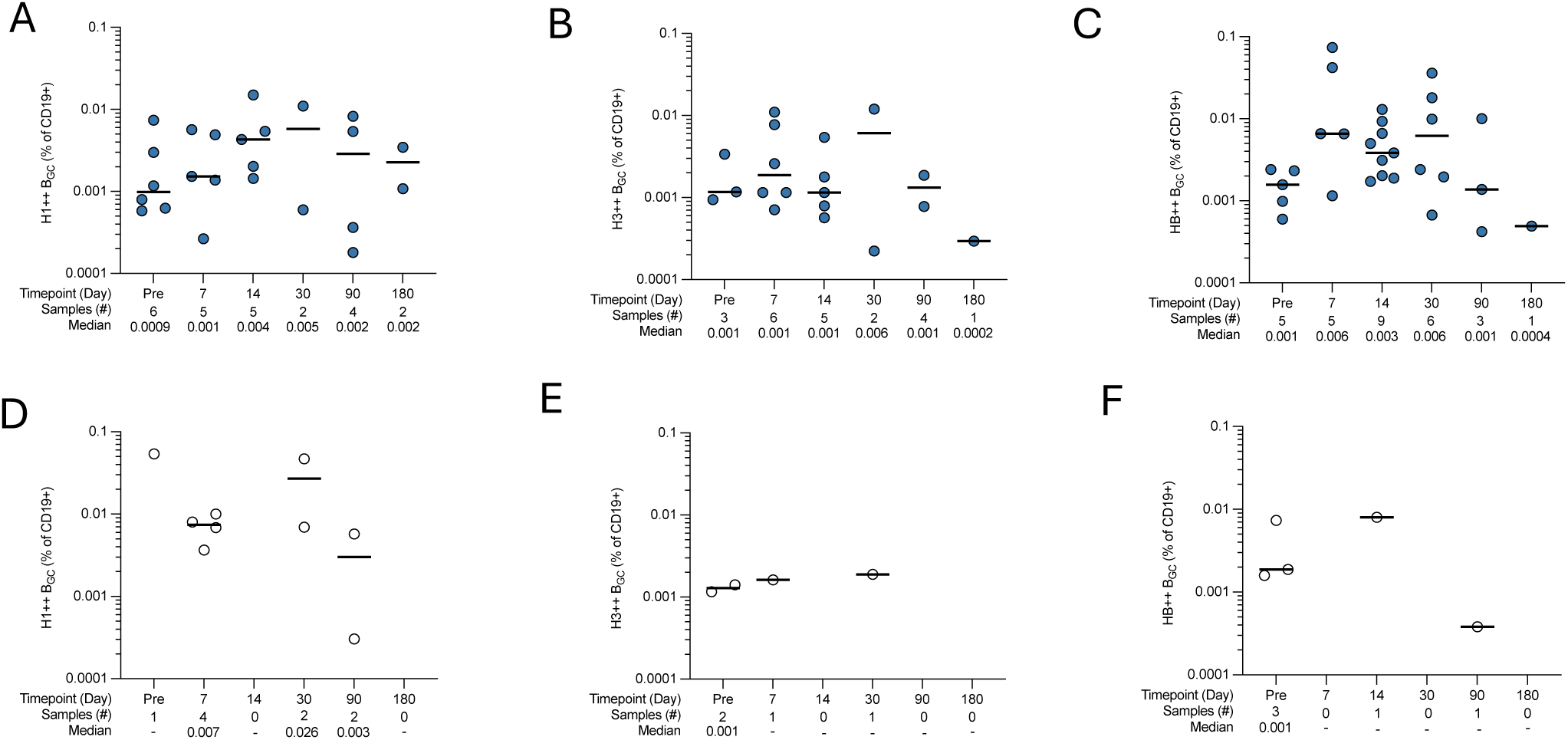
Frequency of antigen specific B_GC_ in NP swabs. **(A-C)** Frequency of antigen specific **(A)** H1-, **(B)** H3-, **(C)** HB- specific B_GC_ in FluMist vaccinees. **(D-F)** Frequency of antigen specific **(D)** H1-, **(E)** H3-, **(F)** HB- specific B_GC_ in IIV recipients. Line represents the median. Only NP swab samples with ≥ 490 naïve (IgD+) B cells and ≥ 3 HA-specific (HA++) B_GC_ cells for the HA-subtype of interest were included in the analysis.

**Fig. S12.**
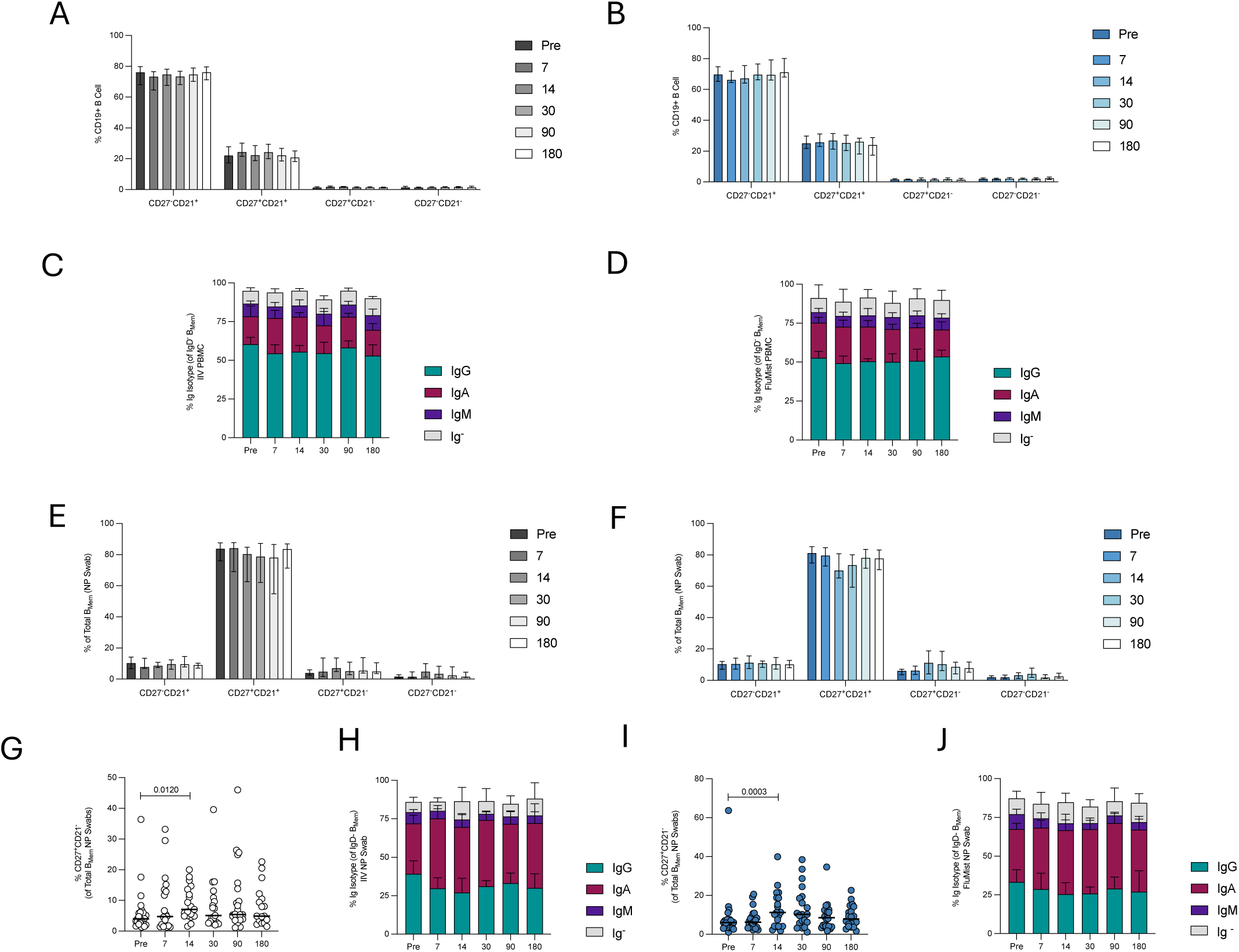
CD27/CD21 and immunoglobulin isotype profiling of non-antigen specific B_Mem_. Frequency of CD21^+/-^/CD27^+/-^ phenotype as % of CD19+ B cells in circulation following **(A)** IIV and **(B)** FluMist. Percentage of IgG, IgA, IgM or Ig- B_Mem_ as % of Total B_Mem_ in circulation in the **(C)** IIV and **(D)** FluMist cohort. **(E-F)** Frequency of CD21^+/-^/CD27^+/-^ phenotype as % of Total B_Mem_ cells in the upper airway (NP) in the **(E)** IIV and **(F)** FluMist cohort. **(G)** Frequency of CD27^+^/CD21^-^ B_Mem_ (% of Total B_Mem_) in the upper airway of IIV recipients. **(H)** Percentage of IgG^+^, IgA^+^, IgM^+^ or Ig^-^ B_Mem_ as % of Total B_Mem_ in the upper airway of IIV recipients. **(I)** Frequency of CD27^+^/CD21^-^ B_Mem_ (% of Total B_Mem_) in the upper airway of FluMist recipients. **(J)** Percentage of IgG^+^, IgA^+^, IgM^+^ or Ig^-^ B_Mem_ as % of Total B_Mem_ in the upper airway of FluMist recipients. **(A-F, H,J)** Data are displayed as median ± 95% confidence intervals. **(G,J)** Data are displayed as the median.

**Fig. S13.**
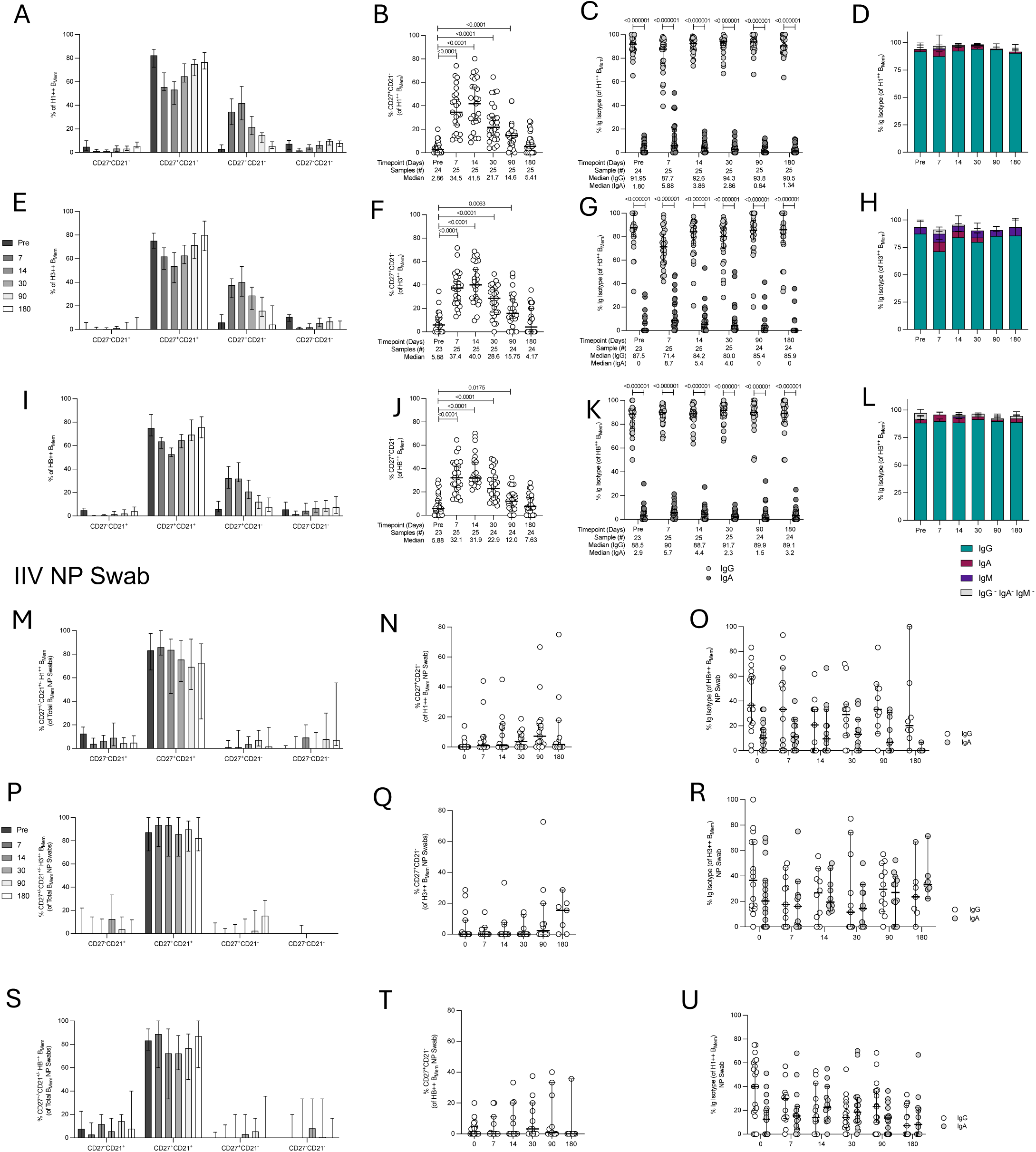
HA subtype-specific characterization of CD27^+^CD21^-^ phenotype and B_Mem_ isotypes in circulation and the upper airway following IIV administration. **(A-L)** Circulating HA subtype specific B_Mem_ frequencies from circulation. **(A)** CD21^+/-^/CD27^+/-^ phenotype H1 B_Mem_ (% of H1^++^ B_Mem_), **(B)** CD27^+^/CD21^-^H1^++^ B_Mem_ (% of H1^++^ B_Mem_), **(C)** IgG^+^ and IgA^+^ H1^++^ B_Mem_, (D) IgG^+^, IgA^+^, IgM^+^ or Ig^-^ B_Mem_ as % of H1^++^ B_Mem_, **(E)** CD21^+/-^/CD27^+/-^ phenotype H3 B_Mem_ (% of H3^++^ B_Mem_), **(F)** CD27^+^/CD21^-^ H3^++^ B_Mem_ (% of H3^++^ B_Mem_), **(G)** IgG^+^ and IgA^+^ H3^++^ B_Mem_, (H) IgG^+^, IgA^+^, IgM^+^ or Ig^-^ B_Mem_ as % of H3^++^ B_Mem_, **(I)** CD21^+/-^/CD27^+/-^ phenotype HB B_Mem_ (% of HB^++^ B_Mem_), **(J)** CD27^+^/CD21^-^ HB^++^ B_Mem_ (% of HB^++^ B_Mem_), **(K)** IgG^+^ and IgA^+^ HB^++^ B_Mem_, **(L)** IgG^+^, IgA^+^, IgM^+^ or Ig^-^ B_Mem_ as % of HB^++^ B_Mem_. (M-U) NP swab HA subtype specific BMem frequencies. **(M)** CD21^+/-^/CD27^+/-^ phenotype H1 B_Mem_ (% of H1^++^ B_Mem_), **(N)** CD27^+^/CD21^-^ H1^++^ B_Mem_ (% of H1^++^ B_Mem_), **(O)** IgG^+^ and IgA^+^ H1^++^ B_Mem_, **(P)** CD21^+/-^/CD27^+/-^ phenotype H3 B_Mem_ (% of H3^++^ B_Mem_), **(Q)** CD27^+^/CD21^-^ H3^++^ B_Mem_ (% of H3^++^ B_Mem_), **(R)** IgG^+^ and IgA^+^ H3^++^ B_Mem_, **(S)** CD21^+/-^/CD27^+/-^ phenotype HB B_Mem_ (% of HB^++^ B_Mem_), **(T)** CD27^+^/CD21^-^ HB^++^ B_Mem_ (% of HB^++^ B_Mem_), **(U)** IgG^+^ and IgA^+^ HB^++^ B_Mem_. **(B,F,J)** Statistical significance between timepoints was assessed by Wilcoxon matched pairs signed rank test. **(C,G,K)** Statistical significance between IgG and IgA was assessed by multiple Wilcoxon matched pairs signed rank test with Holm-Šídák’s multiple comparisons test. Data are displayed as median ± 95% confidence intervals.

**Fig. S14.**
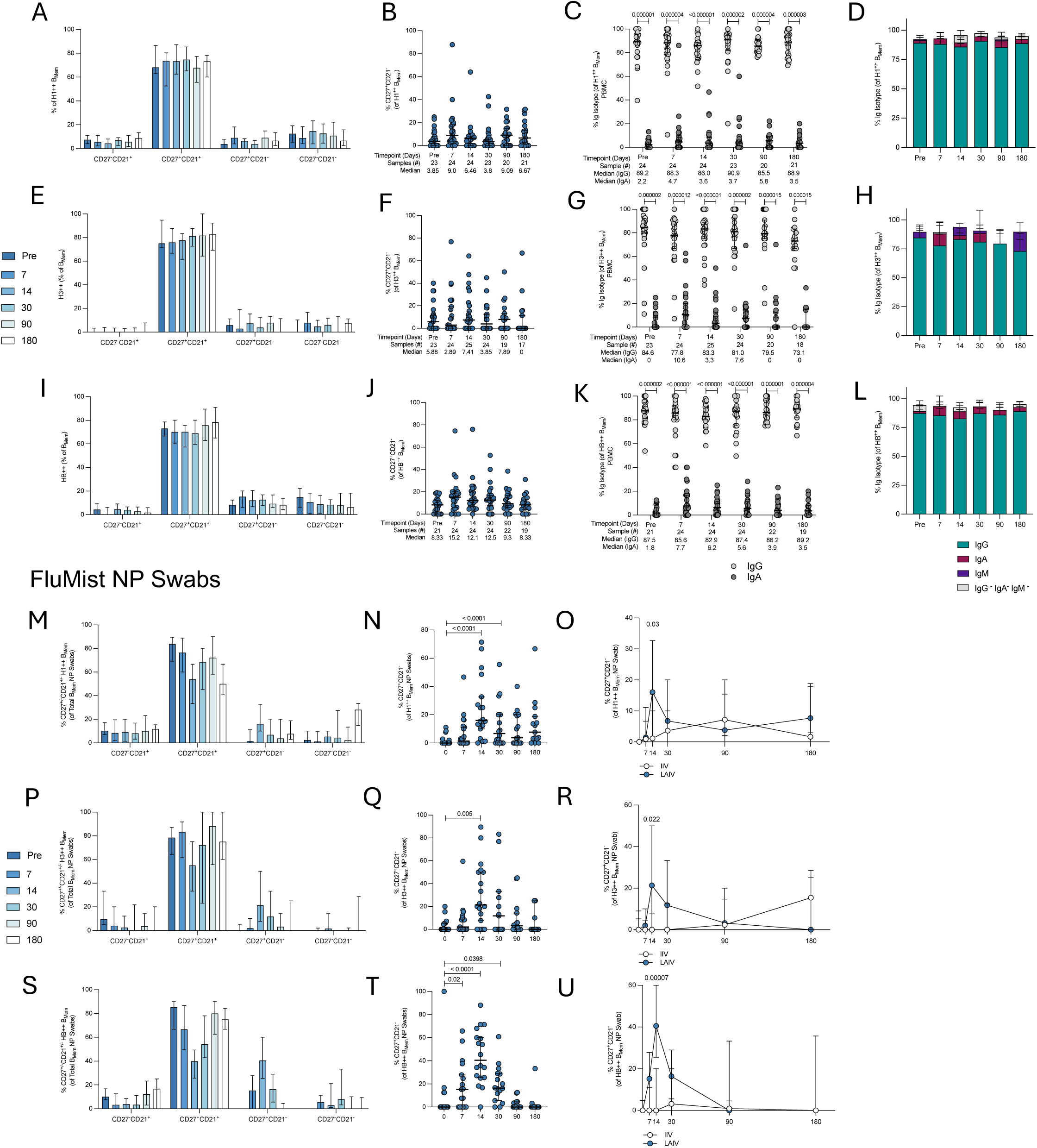
HA subtype-specific characterization of CD27^+^CD21^-^ phenotype and B_Mem_ isotypes in PBMC and NP swabs following FluMist. HA subtype specific frequencies from **(A-L)** PBMC and **(M-U)** NP swabs. Frequency of **(A)** CD21^+/-^/CD27^+/-^ phenotype H1 B_Mem_ (% of H1^++^ B_Mem_), **(B)** CD27^+^/CD21^-^ H1^++^ B_Mem_ (% of H1^++^ B_Mem_), **(C)** IgG^+^ and IgA^+^ H1^++^ B_Mem_, **(D)** IgG^+^, IgA^+^, IgM^+^ or Ig^-^ B_Mem_ as % of H1^++^ B_Mem_, **(E)** CD21^+/-^/CD27^+/-^ phenotype H3 B_Mem_ (% of H3^++^ B_Mem_), **(F)** CD27^+^/CD21^-^ H3^++^ B_Mem_ (% of H3^++^ B_Mem_), **(G)** IgG^+^ and IgA^+^ H3^++^ B_Mem_, **(H)** IgG^+^, IgA^+^, IgM^+^ or Ig^-^ B_Mem_ as % of H3^++^ B_Mem_, **(I)** CD21^+/-^/CD27^+/-^ phenotype HB B_Mem_ (% of HB^++^ B_Mem_), **(J)** CD27^+^/CD21^-^ HB^++^ B_Mem_ (% of HB^++^ B_Mem_), **(K)** IgG^+^ and IgA^+^ HB^++^ B_Mem_, **(L)** IgG^+^, IgA^+^, IgM^+^ or Ig^-^ B_Mem_ as % of HB^++^ B_Mem_; **(M)** CD21^+/-^/CD27^+/-^ phenotype H1 B_Mem_ (% of H1^++^ B_Mem_), **(N)** CD27^+^/CD21^-^ H1^++^ B_Mem_ (% of H1^++^ B_Mem_), **(P)** CD21^+/-^/CD27^+/-^ phenotype H3 B_Mem_ (% of H3^++^ B_Mem_), **(Q)** CD27^+^/CD21^-^ H3^++^ B_Mem_ (% of H3^++^ B_Mem_), **(S)** CD21^+/-^/CD27^+/-^ phenotype HB B_Mem_ (% of HB^++^ B_Mem_), **(T)** CD27^+^/CD21^-^ HB^++^ B_Mem_ (% of HB^++^ B_Mem_). **(O, R, U)** Comparison of **(O)** H1^++^ CD27^+^/CD21^-^, **(R)** H3^++^ CD27^+^/CD21^-^, **(U)** HB^++^ CD27^+^/CD21^-^ B_Mem_ in NP swabs from IIV and FluMist recipients over time following vaccination. **(C,G,K)** Statistical significance between IgG and IgA was assessed by multiple Wilcoxon matched pairs signed rank test with Holm-Šídák’s multiple comparisons test. Data are displayed as median ± 95% confidence intervals.

**Table S1.** Flow Panel for Analysis of PBMC and NP Swabs.2.

| Antibody | Clone | Source | Cat # | Dilution |
| --- | --- | --- | --- | --- |
| Streptavidin- AlexaFluor647 | - | BioLegend | 405237 |  |
| Streptavidin- BV711 (H1) | - | BD Biosciences | 563262 |  |
| Streptavidin- BV786 (H3) | - | BD Biosciences | 563858 |  |
| Streptavidin- BV605 (Influenza B HA) | - | BD Biosciences | 563260 |  |
| Live/Dead Fixable Blue Stain | - | ThermoFisher | L34962 | 1:500 |
| Mouse anti-human CD45 BUV496 | H130 | BD Biosciences | 569101 | 1:500 |
| Mouse anti-human CD19 BUV593 | SJ25C | BD Biosciences | 612916 | 1:500 |
| Mouse anti-human CD16 BV510 (Dump) | 3G8 | BioLegend | 302048 | 1:500 |
| Mouse anti-human CD14 BV510 (Dump) | 63D3 | BioLegend | 367124 | 1:500 |
| Mouse anti-human CD56 BV510 (Dump) | HCD56 | BioLegend | 318340 | 1:500 |
| Mouse anti-human PD-1 BV650 | EH12.2H7 | BioLegend | 329950 | 1:500 |
| Mouse anti-human CD4 cFluor548 | SK3 | Cytek Biosciences | R7-20043 | 1:500 |
| Mouse anti-human CD45RA RB780 | HI100 | BD Biosciences | 569082 | 1:500 |
| Mouse anti-human CD8 cFluorBYG710 | SK1 | Cytek Biosciences | R7-20219 | 1:500 |
| Mouse anti-human CD3 APC | SK7 | BioLegend | 344812 | 1:500 |
| Mouse anti-human CD20 BUV395 | 2H7 | BD Biosciences | 563782 | 1:200 |
| Mouse anti-human IgD Pacific Blue | IA6-2 | BioLegend | 348224 | 1:200 |
| Mouse anti-human IgA PE | IS11-8E10 | Miltenyi Biotech | 130-113-476 | 1:200 |
| Mouse anti-human CD21 PE-Dazzle594 | Bu32 | BioLegend | 354922 | 1:200 |
| Mouse anti-human CD69 PE-Fire640 | FN50 | BioLegend | 310960 | 1:200 |
| Mouse anti-human CD27 Pe-Cy7 | M-T271 | BioLegend | 356412 | 1:200 |
| Mouse anti-human CXCR5 APC-Fire750 | J252D4 | BioLegend | 356946 | 1:200 |
| Mouse anti-human CD38 APC-Fire810 | HB-7 | BioLegend | 356644 | 1:200 |
| Mouse anti-human CCR7 Pe-Cy5 | G043H7 | BioLegend | 353272 | 1:100 |
| Mouse anti-human IgM BV570 | MHM-88 | BioLegend | 314518 | 1:100 |
| Mouse anti-human CD103 BB700 | Ber-ACT8 | BD Biosciences | 745919 | 1:100 |
| Mouse anti-human IgG Alexa Fluor- 700 | G18-145 | BD Biosciences | 561296 | 1:100 |

